# Genomic Determinants of Lethality and Therapeutic Vulnerability in Castration-Resistant Prostate Cancer

**DOI:** 10.64898/2026.06.18.733209

**Authors:** Ryan J. Rebernick, Liat Hammer, Mahnoor Gondal, Abhijit Parolia, Yi-Mi Wu, Noshad Hosseini, Matthew Schipper, Anbarasu Kumaraswamy, Matthew McFarlane, Xuhong Cao, Phillip Palmbos, Megan Caram, David Smith, Sarah Yentz, MI-OncoSeq Team, Stand-Up To Cancer Team, Maha H. Hussain, Thomas C. Westbrook, Daniel Spratt, William C. Jackson, Joshi J. Alumkal, Zachery Reichert, Dan Robinson, Arul M. Chinnaiyan, Robert Dess, Marcin Cieslik

**Author notes:** Corresponding author CORRESPONDING AUTHORS Marcin Cieslik, PhD Robert Dess, MD Arul Chinnaiyan, MD, PhD. These authors contributed equally (Ryan J. Rebernick, Liat Hammer).

## Abstract

Despite extensive characterization, the molecular determinants that shape disease progression and therapeutic outcomes in metastatic castration-resistant prostate cancer (mCRPC) remain incompletely defined. This large, multi-institutional analysis reveals that the genomic determinants of outcomes in primary prostate cancer are distinct from those in castration-resistant disease. We identify androgen-regulated *MYC* fusions as potent primary genetic drivers catalyzing early progression. Additionally, across independent mCRPC cohorts, we validate *TP53, RB1*, and *CDKN1B* alterations as the strongest genetic drivers of poor survival. Mechanistically, *CDKN1B* loss induces an AR-positive epithelial–mesenchymal transition rather than neuroendocrine differentiation, while *TP53* loss triggers whole-genome duplication and subsequent adaptive copy-number gains. Within TP53-altered tumors, *AR* amplifications confer improved survival while *AR* mutations are protective regardless of *TP53* status. Integrating these genomic events with transcriptomic phenotypes, we derive CAPrisk, a multi-omic prognostic classifier. CAPrisk stratifies patients with mCRPC into three risk groups with >33-month survival differences and predicts outcomes across androgen receptor pathway inhibitors, taxanes, PARP-inhibitors, and radium-223. Together, these findings expand the clinical utility of genomic profiling in advanced prostate cancer and establish a framework for risk stratification in lethal disease.

**IN BRIEF (eTOC BLURB):** Rebernick et al. analyze a multi-institutional cohort of 1,331 prostate cancer samples and identify the genomic determinants of progression to lethal, castration-resistant disease. They define rare AR-regulated MYC fusions and haploinsufficient CDKN1B loss as drivers of poor outcome while highlighting AR alterations as predictors of improved prognosis. They develop CAPrisk, a multi-omic classifier that stratifies patient outcomes across androgen receptor pathway inhibitors, taxanes, PARP-inhibitors, and radium-223.

**HIGHLIGHTS:**

– AR-regulated *MYC* fusions are rare (∼2%) truncal events driving aggressive disease.
– Distinct *AR* alterations drive favorable outcomes in *TP53*-altered and non-altered contexts.
– Haploinsufficient *CDKN1B* loss predicts poor survival in AR-positive tumors.
– A multi-omic model predicts outcomes across four standard-of-care therapies.

**MI-OncoSeq Team:** Chandan Kumar, Erica Rabban, Kayla Muschong, Lakshmi P. Kunju, Javed Siddiqui, Yu Ning, Rui Wang, Fengyun Su, Yelena Kleyman-Smith, Josh N. Vo, Jin Chen, Rahul Mannan

**Stand Up To Cancer Team:** Wassim Abida, Joanna Cyrta, Glenn Heller, Davide Prandi, Joshua Armenia, Ilsa Coleman, Matteo Benelli, Eliezer M. Van Allen, Andrea Sboner, Tarcisio Fedrizzi, Juan Miguel Mosquera, Brian D. Robinson, Navonil De Sarkar, Lakshmi P. Kunju, Scott Tomlins, Daniel Nava Rodrigues, Massimo Loda, Anuradha Gopalan, Victor E. Reuter, Colin C. Pritchard, Joaquin Mateo, Diletta Bianchini, Susana Miranda, Suzanne Carreira, Pasquale Rescigno, Julie Filipenko, Jacob Vinson, Robert B. Montgomery, Himisha Beltran, Elisabeth I. Heath, Howard I. Scher, Philip W. Kantoff, Mary-Ellen Taplin, Nikolaus Schultz, Johann S. deBono, Francesca Demichelis, Peter S. Nelson, Mark A. Rubin, Charles Sawyers

## INTRODUCTION

Prostate cancer (PCa) is the most common malignancy in adult men, estimated to account for 31% of new cancer diagnoses in 2026 ^1^. Most cases are localized at diagnosis and can be managed with curative intent. However, 20-30% of patients will either present with or later develop metastatic castration-sensitive prostate cancer (mCSPC) ^2^. Despite advances in combination systemic therapy, most patients with mCSPC will eventually develop resistance to hormonal therapy, progress to metastatic castration-resistant prostate cancer (mCRPC), and die of their disease and its complications ^3–7^.

Large-scale sequencing initiatives have comprehensively mapped the genomic and transcriptomic landscape of prostate cancer. This has led to the discovery of multiple primary genetic drivers including mutually exclusive truncal ETS and RAF gene fusions ^8–10^, as well as mutations of *SPOP* ^10,11^*, FOXA1* ^12^, and *CDK12* ^13^. In addition, while mutations in homologous recombination (HR) and mismatch repair (MMR) genes ^14,15^ are not strictly mutually exclusive, these alterations are often grouped within a truncal class as they are frequently identified in localized disease and can occur through germline alterations ^14–16^.

Subsequent studies have identified recurrent alterations in advanced and metastatic prostate cancer, particularly in key genes such as *AR*, *PTEN*, *TP53*, and *RB1* ^17–19^. Furthermore, co-alterations in *TP53* and *RB1* are heavily enriched in neuroendocrine prostate cancer, an AR-independent phenotype which develops in response to inhibition of androgen receptor signaling ^20–22^. Prominent transcriptomic subtypes of mCRPC include luminal and basal-like lineages, highly proliferative variants, and those enriched for WNT, TGF-β, and mTOR signaling ^23–26^. While localized prostate cancers have relatively stable genomes and low tumor mutation burden (TMB), chromosomal instability (CIN) and TMB are markedly increased in metastatic tumors ^27^.

Despite the extensive characterization of both the primary and metastatic prostate cancer genomic landscape ^10,17,18,28–30^, current translational knowledge is skewed toward primary disease. Similarly, prognostic classifiers (e.g. Decipher, GPS, Prolaris) are primarily implemented in localized, hormone-naive settings ^31–38^. This progress underscores a disparity in the management of lethal disease, as the genomic and transcriptomic architectures that dictate clinical outcomes in mCRPC remain incompletely defined.

Within mCRPC, alterations in *TP53*, *AR*, and *RB1* are associated with shorter time on treatment with androgen receptor pathway inhibitors (ARPIs) and *RB1* alterations linked with decreased overall survival (OS) from time of biopsy ^19,39–41^. Shorter OS in mCRPC is also associated with select transcriptomic phenotypes, including basal and neuroendocrine signatures, loss of androgen signaling, and elevated proliferative and mTOR activity ^23,25,42,43^. Studies of liquid biopsies from patients with mCRPC have reported additional associations between *AR* splice variants and *TP53* alterations with OS from the time of biopsy ^44,45^. However, the mechanistic trajectory by which *TP53* inactivation drives ARPI resistance remains largely unexplored, particularly the extent to which these distinct, instability-driven *AR* alterations dictate differential resistance patterns to standard-of-care inhibitors. Additionally, these circulating tumor DNA analyses have linked alterations in *TP53* and HR genes (*BRCA2*, *ATM*) to shorter time to progression on ARPIs ^45^. Conversely, no association between HR-deficiency (HRd) status and time on ARPIs was found in solid tumor profiling ^19^.

Here, we present *CArcinoma of Prostate Sequencing of TumOr and cliNical Endpoints* (CAPSTONE), one of the largest, clinico-genomic compendia of lethal prostate cancer to date. Leveraging a comprehensive, uniform computational analysis of 1,331 samples from 1,264 patients, we dissect the impact of genetic alterations and transcriptional phenotypes on clinical outcomes in primary PCa and mCRPC. We demonstrate that the genomic drivers predicting lethal potential in primary hormone-sensitive disease are distinct from the alterations that dictate survival and treatment resistance in the metastatic castration-resistant setting. We translate these findings into a prognostic tool for mCRPC, termed CAPrisk, which stratifies patient risk and predicts outcomes following standard-of-care therapies.

## RESULTS

### Genomic integration of 1,331 samples from 1,264 prostate cancer patients

To systematically define the genomic and transcriptomic determinants of clinical outcomes across the full spectrum of disease progression, we established CAPSTONE, an mCRPC-enriched cohort that integrates rigorous clinical follow-up with simultaneous DNA and RNA characterization. CAPSTONE comprises 1,331 unique prostate cancer samples obtained from 1,264 patients derived from 3 distinct sources - internal University of Michigan (U-M) MI-OncoSeq ^12,13,29,46^ patients (*Discovery*), patients from six additional institutions enrolled through the Stand Up To Cancer (*SU2C*) International Dream Team, and patients profiled through The Cancer Genome Atlas (TCGA).

For the *Discovery* cohort we identified a total of 404 U-M patients diagnosed with prostate cancer between January, 1980 and May, 2021. Of these 404 patients, 324 (80.2%) had clinical data and at least one tumor sample sufficient for sequencing (**Figure 1A**). In total, 352 independent prostate cancer samples were analyzed. The majority (232/352, 66%) were obtained from patients with mCRPC. The cohort also included tumors from patients with recurrent mCSPC (47/352, 13%), de novo mCSPC (38/352, 11%), and localized prostate cancer (35/352, 10%). In total, the *Discovery* cohort consisted of 324 patients profiled across 352 unique samples.

**Figure 1:**
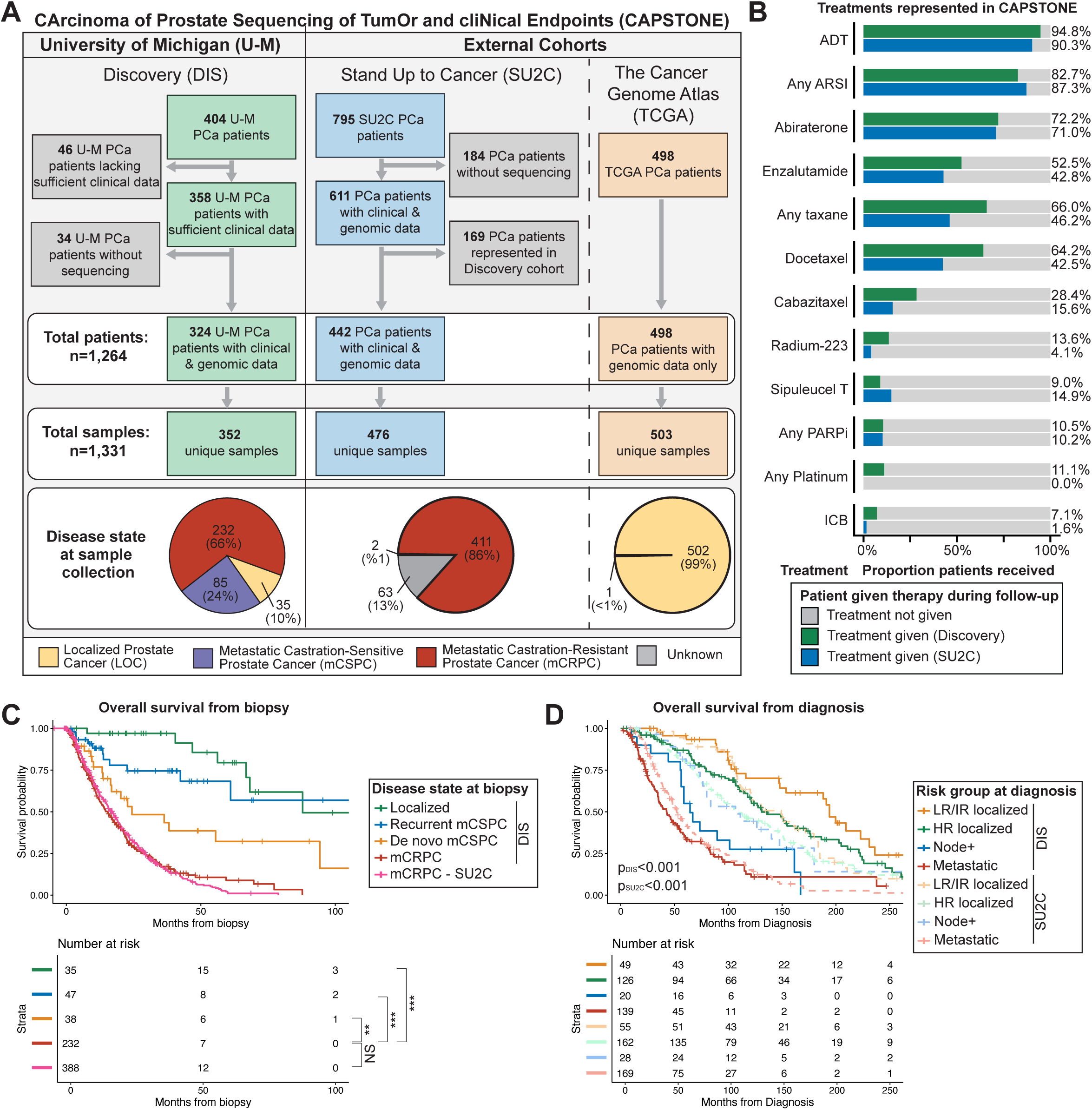
A multi-institutional clinico-genomic cohort of 1,331 prostate cancer samples from 1,264 patients. **A.** CAPSTONE samples and patients stratified by cohort and disease state at time of sample collection. **B.** Treatment information for *Discovery* and Stand Up To Cancer (SU2C) cohorts. Bars indicate the proportion of patients that received the therapy during follow up. **C.** Overall survival from time of biopsy stratified by disease state at biopsy and cohort. Abbreviations: mCSPC (metastatic castration-sensitive prostate cancer), mCRPC (metastatic castration-resistant prostate cancer). Significance: * <0.05, ** <0.01, ***<0.001. **D.** Overall survival from the time of diagnosis stratified by risk group at diagnosis for the Discovery and SU2C cohorts. P-value represents a log-rank test for Discovery and SU2C cohorts respectively.

The SU2C cohort included patients from 7 member institutions ^19^. To ensure this dataset was fully independent of the *Discovery* cohort, we excluded all U-M patients that were represented in the *Discovery* cohort. This left 442 (56%) remaining patients with available sequencing (**Figure 1A**). From these 442 patients, 476 independent tumor samples were available. The majority of these samples (411/476, 86%) were obtained from mCRPC, while 2 (1%) were from recurrent mCSPC, and 63 (13%) had an undetermined disease state at time of metastatic sample collection. In total, the SU2C cohort contained 442 patients profiled across 476 samples. Lastly, the TCGA cohort encompassed 498 patients with prostate cancer, with the majority of samples (502/503, 99%) obtained from localized prostate cancer (**Figure 1A**).

For each sample across the *Discovery*, SU2C, and TCGA cohorts, raw DNA and RNA sequencing data was retrieved and uniformly processed using a comprehensive genomics pipeline (**Methods**). All 1,331 samples were reviewed for sufficient sequencing quality (**Table S1**). Across the *Discovery* (n=294), SU2C (n=295), and TCGA (n=443) cohorts, 1,032 samples (76%) from 1,003 patients (79%) had high-quality RNA and DNA sequencing as defined by strict QC parameters (**Methods**), including tumor purity. Metastatic samples originated from diverse sites, including prostate, bone, liver, and soft tissue (**Table S2**). Standardized bioinformatics methods were applied across all samples, as previously described ^29,47^, to perform somatic and germline variant calling, identify gene fusions, and quantify copy number alterations, including loss-of-heterozygosity. We used a systematic approach to define allelic status, characterize genetic drivers, and annotate disease-specific pathways (**Methods**).

### Clinical characterization of the *Discovery* and SU2C cohorts

Clinical records in the CAPSTONE Discovery cohort were comprehensively curated to capture TNM staging, International Society of Urological Pathology grade groups, prostate-specific antigen levels, and therapeutic history, alongside critical endpoints including biochemical recurrence, metastasis, castration-resistance, and overall survival (**Methods**). Similarly, all available clinical information for the SU2C cohort was systematically curated ^19^.

The Discovery and SU2C cohorts were demographically and clinically comparable, featuring extensive longitudinal follow-up from the time of diagnosis (DIS:68.1, SU2C:80.1 months) (**Table S3**). The median follow-up time after sample collection across all disease states was 15.4 months (IQR: 6.9-30.0) for Discovery and 13.9 months (IQR: 6.1-25.3) for SU2C samples. Central histopathologic review identified neuroendocrine histology in 9 of 352 (3%) Discovery samples, while neuroendocrine histology was detected in 12 of 476 (3%) SU2C samples. Validating our clinical data, OS from sample collection was significantly shorter for mCRPC compared to de novo mCSPC (p=0.0014), recurrent mCSPC (p<0.001), and localized disease (p<0.001) (**Figure 1C**). OS from diagnosis was also significantly associated with risk groups at diagnosis in both the *Discovery* (p<0.001) and *SU2C* (p<0.001) cohorts (**Figure 1D**). There was no difference in OS from sample collection between *Discovery* and SU2C mCRPC samples (p=0.78).

Both the *Discovery* and SU2C cohorts contained detailed treatment information. A high proportion of patients received standard-of-care therapies, including ADT (94.8% *Discovery*, 90.3% SU2C), ARPIs (82.7% *Discovery*, 87.3% SU2C), and taxane chemotherapy (66.0% *Discovery*, 46.2% SU2C) (**Figure 1B**). Additionally, some patients received PARP inhibitors (PARPi), Sipuleucel-T, platinum chemotherapy, Radium-223, and immune checkpoint blockade (ICB). Given that most tumor samples were from mCRPC, the majority had received ADT (79.0%, 88.0%) and ARPIs (50.3%, 65.8%) prior to sample collection (**Figure S1A**).

### The progressive genomic landscape of advanced prostate cancer

We utilized 1,008 high-quality samples with known disease states at time of sample collection to define the landscape of recurrent genomic alterations across prostate cancer disease states ^14,17–19,29^. By simultaneously profiling germline and somatic mutations, copy number alterations, and gene fusions, we systematically evaluated the status of known drivers mapped onto signaling pathways.

Comparing samples from mCRPC to those from localized disease, we observed expected increases in *AR* (54% vs 1%), *TP53* (55% vs 21%), and *PTEN* (41% vs 18%) alterations (**Figure 2A**). The shift from localized disease to mCSPC was marked by higher alteration rates in APC ^48^, as well as *TP53*, *RB1*, BRCA2, and *PTEN*. In contrast, the subsequent progression from mCSPC to mCRPC was only marked by an increase in *AR* alterations (**Figure 2A**). Somatic HRd alteration rates nominally increased from mHSPC to mCRPC (4% vs 13%, p=0.053), with total rates of HRd (somatic and germline) peaking at 18% in mCRPC (**Figure S1B-C**). Alterations in PI3K and WNT pathways increased primarily from localized to mCSPC samples (17% vs 43%, p=6.5e-7; 4% vs 21%, p=2.1e-7 respectively), but were altered at comparable rates between mCSPC and mCRPC samples (43% vs 38%, p=0.49; 21% vs 13%, p=0.13 respectively) (**Figure S1D-E**).

**Figure 2:**
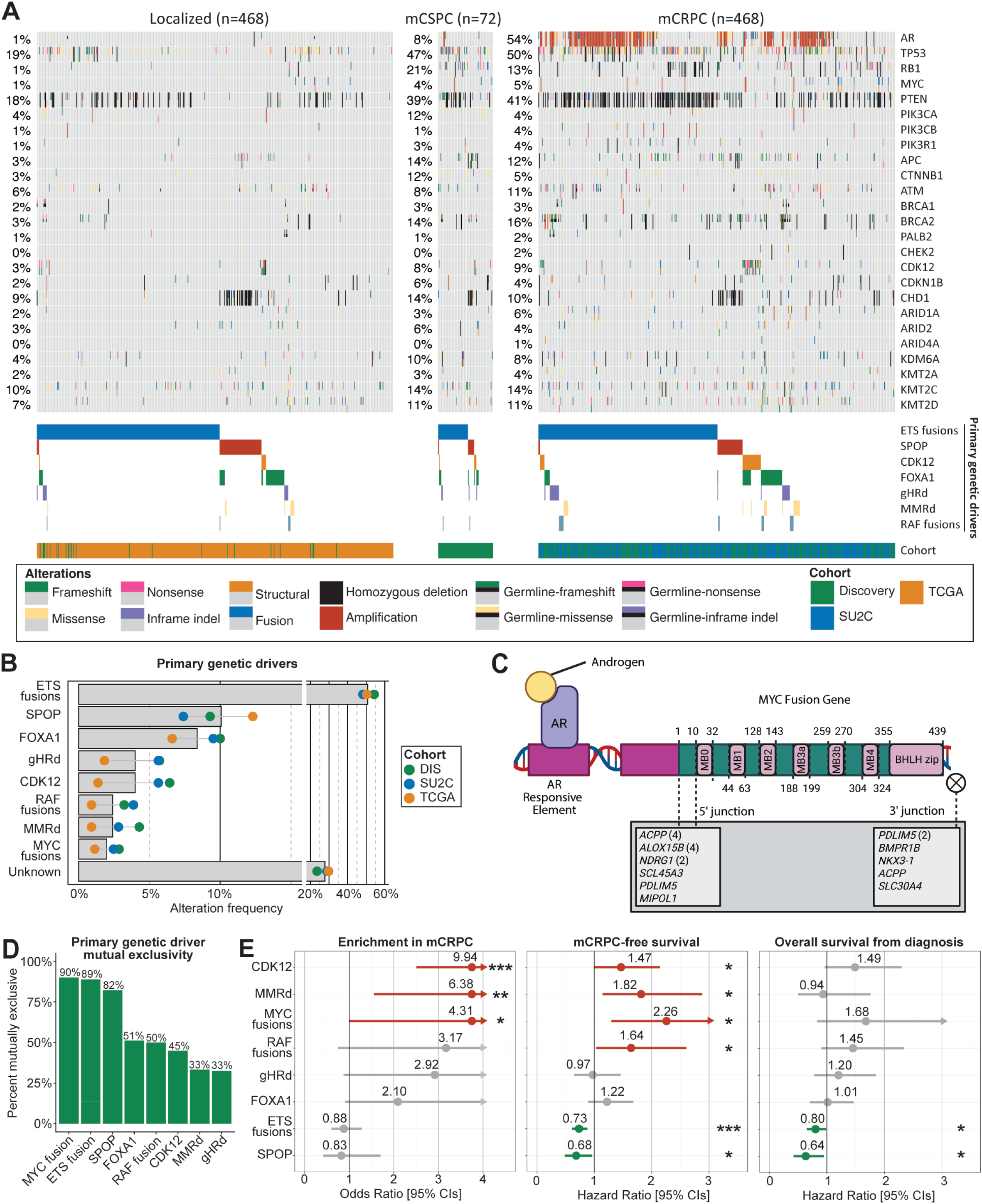
Primary genetic drivers and their role in lethal prostate cancer. **A.** Oncoprint of 1,008 high-quality prostate cancer samples with known time of sample collection. Genomic alterations include somatic and germline mutations, somatic focal amplifications and homozygous deletions, and curated primary genetic driver annotations. **B.** The frequency of established primary genetic drivers in prostate cancer across the 3 CAPSTONE component cohorts. Abbreviations: germline homologous recombination deficiency (gHRD), mismatch repair deficient (MMRd). **C.** Identified MYC fusions and their interaction with the Androgen Receptor (AR). MYC protein domains shown with amino acid positions. Fusion partner genes listed with the number of unique patients detected in shown in parentheses. **D.** Mutual exclusivity of primary genetic drivers within CAPSTONE. **E.** Relationship between primary genetic drivers and enrichment in cases of lethal prostate cancer as compared with benign prostate cancer as well as their relationship with metastatic castration-resistant prostate cancer free survival (mCRPCfs) and overall survival from diagnosis. Point estimates represent either odds ratios or hazard ratios. Lines represent 95% confidence intervals. Arrowheads indicate the interval extends beyond range shown. Significance: * <0.05, ** <0.01, ***<0.001.

We next shifted our focus from specific genetic drivers to global measures of chromosomal instability (CIN). We hypothesized that the selective pressure from androgen deprivation would drive a continuous increase of chromosomal instability ^49^, yet we observed a complex dichotomy. While standard instability metrics (nLOH, nLST) rose in mCSPC, they plateaued upon progression to mCRPC, suggesting a saturation of large-scale structural changes (**Figure S1F-G**). Conversely, evidence of ongoing evolution persisted in the form of whole genome duplication, focal gains, the weighted genome integrity index (wGII), and number of telomeric allelic imbalances (nTAI), which accumulated in a stepwise fashion throughout the disease course (**Figure S1H-K**). Notably, this sustained telomeric allelic imbalance appears to be the primary driver of high HRd scores ^50,51^ in the mCRPC setting, as other components of HRd signatures stabilize (**Figure S1L**).

### MYC gene fusions are primary genetic drivers of prostate cancer

Primary genetic drivers are recurrent, mutually exclusive truncal alterations that arise early in prostate cancer and define distinct molecular subtypes ^8–14^. We analyzed the frequency of primary genetic drivers across CAPSTONE and found alteration rates comparable to prior studies (**Figure 2B**). ETS gene fusions were the most common (510/1003, 51%), followed by *SPOP* hotspot mutations (101/1003, 10%), *FOXA1* class I mutations (84/1003, 8%), germline homologous recombination deficiency (gHRd) (40/1003, 4%), and *CDK12* alterations (40/1003, 4%). MMRd and RAF fusions were comparatively rare (24/1003, 2%, 24/1003, 2%).

Notably, in tumors lacking a known primary driver (281/1003, 28%), we observed recurrent gene fusions involving the *MYC* oncogene (**Figure 2A**) ^28^. *MYC* encodes c-Myc, a transcription factor frequently overexpressed and recurrently amplified in prostate cancer that serves as a canonical oncogene in prostate cancer genetically engineered mouse models ^52–54^. Across CAPSTONE, 20 (2.0%) patients harbored *MYC* gene fusions, including 5 in localized disease with no other known truncal alteration (**Table S4**). Most *MYC* gene fusions involved either a recurrent 5’ locus joined to the promoter, exon 1, or exon 2 of *MYC* or full length *MYC* joined to a downstream regulatory element of another gene (**Figure 2C**). Notably, exon 2 encodes the N-terminal transcriptional activation domain and contains the first of several highly conserved MYC boxes ^55,56^. Thus, *MYC* fusions preserve all functional domains and annotated regulatory motifs in c-Myc.

Recurrent *MYC* fusion partners include the genes *ACP3* (4/20, 20%), *ALOX15B* (4/20, 20%), *PDLIM5* (3/20, 15%), and *NDRG1* (2/20, 10%). Additional partners include well-known ETS-fusion 5’ genes (or their enhancers) such as *SLC45A3*, *MIPOL1,* and *ACP3.* Strikingly, almost all *MYC* fusions, including with the novel partner *BMPR1B*, involve AR-regulated genes (*PDLIM5*, *ACPP, ALOX15B, SLC45A3, ACP3, NDRG1, NKX3-1, SLC30A4*) ^57–61^. Samples with *MYC* fusions exhibited significantly higher *MYC* expression (p=6.7e-3), but comparable gene copy number (p=0.80) to non-fusion samples (**Figure S2A-B**). Relative to samples with focal *MYC* amplifications, *MYC* fusions had significantly lower MYC gene copies (p=1.5e-9) and comparable MYC expression levels (p=0.81) (**Figure S2A-B**).

Importantly, *MYC* fusions were largely mutually exclusive with established primary genetic drivers with only 2 of 20 (10%) samples harboring concurrent driver alterations (**Figure S2C**). This concurrence rate is lower than established drivers like *ETS* fusions (11%) or *SPOP* mutations (18%), suggesting a primary oncogenic role (**Figure 2D**).

We propose that *MYC* fusions represent a distinct biological entity from *MYC* gains which are prevalent in mCRPC. While gains and amplifications preserve the regulation of *MYC* by its promoter and enhancers ^62,63^, fusions subvert androgen-mediated repression to place *MYC* under direct androgen stimulation. Supported by their mutual exclusivity with established drivers, these data identify *MYC* fusions as a primary driver in localized disease.

### Prognostic significance of primary genetic drivers

Despite defining distinct molecular subtypes, the long-term prognostic significance of primary genetic drivers is incompletely established ^64–67^. We sought to determine whether these truncal genomic events merely initiate tumorigenesis or continue to influence clinical outcomes across the progression of lethal prostate cancer. To address this question, we compared the prevalence of primary genetic drivers between lethal and indolent disease as performed in *Pritchard et al. 2016* ^14^. As primary genetic drivers are detected earliest in prostate cancer development, we directly compared the frequency of driver alterations between patients with mCRPC and ISUP Gleason grade ≤2 localized prostate cancer, a population unlikely to develop lethal disease ^68^.

Compared to men with indolent disease, patients with mCRPC had significantly higher odds of harboring *CDK12* alterations (OR: 9.94 [2.50–408], p=5.1e-4), MMRd (OR: 6.38 [1.55–267], p=6.0e-3), or *MYC* fusions (OR: 4.31 [1.00–184], p=0.046), while gHRd was nominally enriched (OR: 2.92 [0.88-15.3], p=0.078) (**Figure 2E**). Notably, none of the patients with indolent prostate cancer had *CDK12* alterations, MMRd, or *MYC* fusions. Alterations in ETS, *SPOP*, *FOXA1*, and RAF occurred at comparable frequencies between mCRPC and indolent disease.

We next evaluated the prognostic impact of primary genetic drivers among patients who develop lethal disease. This approach is robust, as alteration in those genes are truncal and shared across all metastatic lesions ^69^ (**Methods**), specifically assessing mCRPC-free survival (mCRPCfs) and overall survival (OS). In a univariate analysis, *CDK12* (HR: 1.46, 1.00-2.14, p=0.048), MMRd (HR: 1.82, 1.14-2.88, p=0.011), RAF fusions (HR: 1.64, 1.03-2.60, p=0.035), and *MYC* fusions (HR: 2.26, 1.30-3.93, p=2.2e-3) were associated with shorter mCRPCfs (**Figure 2E**). In contrast, both ETS gene fusions (HR: 0.73, 0.61-0.88, p=7.6e-4) and *SPOP* mutations (HR: 0.68, 0.49-0.96, p=0.029) were associated with prolonged mCRPCfs (**Figure 2E**). ETS fusions and *SPOP* mutations were also associated with a prolonged OS from diagnosis (OSdx) (HR: 0.80, 0.65-0.98, p=0.033, HR: 0.63, 0.42-0.95, p=0.025, respectively) (**Figure 2E**).

Despite its strong association with developing lethal disease and shorter mCRPCfs, MMRd was not significantly associated with OSdx (p=0.84) (**Figure 2E**). This finding may be attributable to the use of immune checkpoint blockade (ICB) (7/20, 35%) which has shown promising efficacy in this patient population ^15,70^. Importantly, no primary genetic driver was significantly associated with OS following progression to mCRPC (OSmcrpc) (**Figure S2D**). Thus, primary genetic drivers influence the rate of progression to mCRPC, but do not independently predict survival after the disease becomes castration-resistant.

### Genomic and transcriptomic determinants of prognosis in mCRPC

To define the biological determinants of survival in lethal prostate cancer, we leveraged the CAPSTONE *Discovery* and *SU2C* cohorts for sequential discovery and validation of genes and pathways critical for prognosis in lethal prostate cancer. We integrated clinical outcomes with a systematic screen of exome-wide somatic alterations and nearly 1,400 transcriptional signatures to identify robust predictors of prognosis in mCRPC (**Figure 3A**, **Methods**).

**Figure 3:**
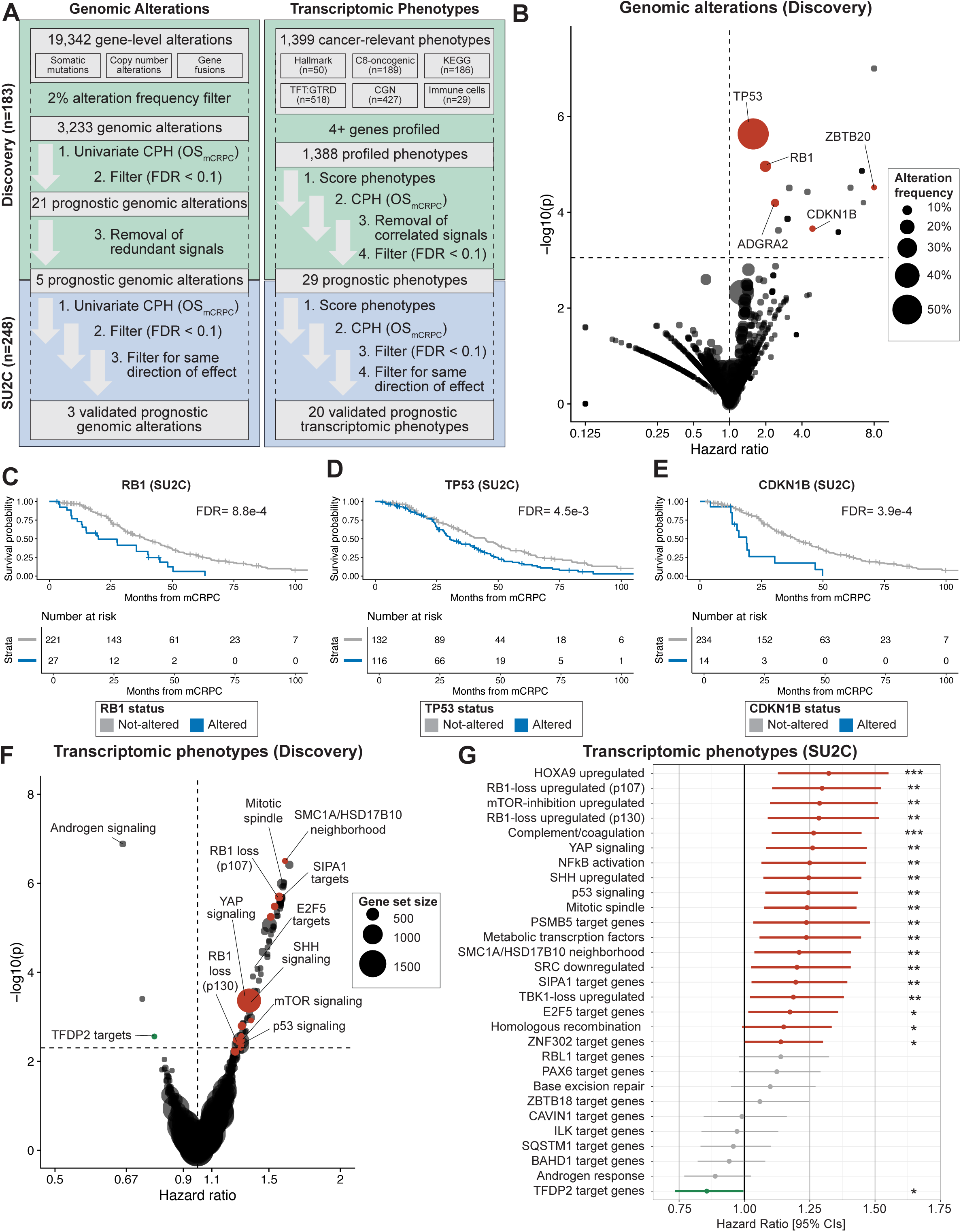
Comprehensive association of genomic alterations and transcriptomic phenotypes with prognosis in lethal prostate cancer. **A.** Overview of the methodology used to identify and validate transcriptomic phenotypes and genomic alterations associated with overall survival from the time of metastatic castration-resistant prostate cancer development (OSmcrpc). Abbreviations: CPH: Cox Proportional Hazards; FDR: False discovery rate. **B.** Volcano plot of hazard ratios and unadjusted p-values from CPH models for 3,233 genomic features in the Discovery cohort. The 5 significant prognostic genomic alterations are labelled. **C-E.** Kaplan meier plot for OSmcrpc in the SU2C cohort for **C)** *RB1*, **D)** *TP53*, and **E)** *CDKN1B* alterations. **F.** Volcano plot of hazard ratios and unadjusted p-values from CPH models for 1,388 transcriptomic phenotypes in the Discovery cohort. The horizontal dotted line represents FDR of 10%. Vertical dotted line corresponds to a hazard ratio of 1. Values to the right of the line are associated with poor prognosis. Dot size represents the alteration frequency. Select transcriptomic phenotypes labelled. **G.** Hazard ratio for 29 significant prognostic transcriptomic phenotypes and OSmcrpc. Point estimates represent hazard ratios, lines represent 95% confidence intervals in the SU2C cohort. Significance: * <0.05, ** <0.01, ***<0.001.

In the *Discovery* cohort, we first evaluated the association of genetic drivers with clinical outcomes in mCRPC. We integrated exome-wide somatic mutations, gene fusions, and copy number alterations to identify specific DNA-level alterations that robustly stratified patient survival (**Figure 3A**). In the *Discovery* cohort, this approach nominated 21 genes significantly (FDR < 0.1) associated with OSmcrpc (**Table S5**). To isolate driver events from passenger copy number effects, we excluded 16 redundant nearby genes mapping to the *RB1* (homozygous deletion) or *ADGRA2* (amplification) loci (**Table S6**). This resulted in five unique prognostic signals: *TP53* (HR^DIS^: 2.30 [1.63-3.25], FDR=3.8e-3), *RB1* (HR^DIS^: 2.98 [1.83-4.84], FDR=7.4e-3), *CDKN1B* (p27/p27KIP1) (HR^DIS^: 5.59 [2.24-13.9], FDR=0.040), *ZBTB20* (HR^DIS^: 9.26 [3.25-26.4], FDR=0.011), and *ADGRA2* (HR^DIS^: 3.51 [1.90-6.51], FDR=0.017) (**Figure 3B**). Upon assessment in the SU2C validation cohort, 3 of these 5 candidates were confirmed as robust drivers of lethal outcomes: *RB1*, *TP53*, and *CDKN1B* (**Figure 3C-E**). *ZBTB20* and *ADGRA2* failed to validate (**Table S5**). Notably, while *PTEN* deletions and *SPOP* mutations are key prognostic factors in primary disease ^66,71^, neither showed an association with survival in the castration-resistant setting. Similarly, collective alterations in pathways including PI3K, HRd, MMRd, or WNT were not significantly associated with survival in mCRPC (**Figure S3A-D**). Only the cell cycle pathway (*RB1*, *CCND1, CDKN2A, CDKN1B*) was associated with OSmcrpc (p=1.3e-9) and this remained significant even after exclusion of *RB1* and *CDKN1B* alterations (p=8.4e-3) (**Figure S3E-F**).

We next applied our dual-cohort approach to systematically discover and validate 1,388 cancer-relevant phenotypes (transcriptomic signatures) with OSmcrpc (**Figure 3F**). We identified 84 significant phenotypes (FDR < 0.1), which were subsequently pruned for redundancy (R>0.7) to define 29 independent prognostic variables. When assessed in the SU2C cohort (validation) , 20 of these 29 independent signals remained significantly associated with OSmcrpc (FDR < 0.1), demonstrating high cross-cohort reproducibility (**Table S7,** **Figure 3G**). The 20 validated prognostic phenotypes revealed a clear biological dichotomy between checkpoint-deficient proliferation and differentiation-induced dormancy. Nineteen signatures were associated with poor survival, predominantly reflecting tumor suppressor loss and unchecked replication. These phenotypes validated the prognostic association of *TP53* and *RB1* genetic loss, while identifying transcriptional programs promoting epithelial-mesenchymal transition (EMT) *i.e.* Sonic hedgehog, Hippo/Yap, and NFkB signaling pathways. In contrast, the sole phenotype associated with improved survival was a signature of *TFDP2* targets, representing a transcriptional program of differentiation and cell cycle exit ^72^. Notably, androgen signaling was markedly associated with prolonged OSmcrpc in the Discovery cohort (HR^DIS^: 0.66, [0.56-0.77], FDR=1.8e-4), but marginally failed to reach significance in the SU2C cohort despite a comparable direction and magnitude of effect (HR^SU2C^: 0.89, [0.77-1.03], FDR=0.13). However, androgen signaling was significant in pooled cohort analysis (HR^comb^: 0.82, [0.74-0.91], FDR=2.8e-3).

Collectively, our findings confirm the established roles of *TP53* and *RB1*, and discover *CDKN1B* as a novel, independent driver of poor prognosis in mCRPC.

### *CDKN1B* loss drives an epithelial mesenchymal transition in lethal prostate cancer

Across both Discovery and SU2C cohorts, the median OSmcrpc for *CDKN1B*-altered patients was 15.8 months, significantly worse than the 35.4 months observed for non-altered patients (p=3.6e-7) (**Figure 4A**). To clarify the basis of this aggressive clinical phenotype, we characterized the spectrum of *CDKN1B* (p27KIP1) alterations across CAPSTONE. In aggregate, 3.4% (36/1032) of the high-quality samples harbored somatic alterations in *CDKN1B* (**Figure 4B**, **Table S8**). Mutations represented nearly half (17/36, 47%) of *CDKN1B* alterations, manifesting as predominantly clonal (13/17, 76%), inactivating events that abolish p27KIP1 protein function through early truncation (17/17, 100%) (**Figure 4C**). Focal homozygous deletions constituted the second most common alteration mechanism (14/36, 39%), complemented by diverse structural variants (5/36, 14%) that included start-loss events and truncating gene fusions (**Figure 4D-E**).

**Figure 4:**
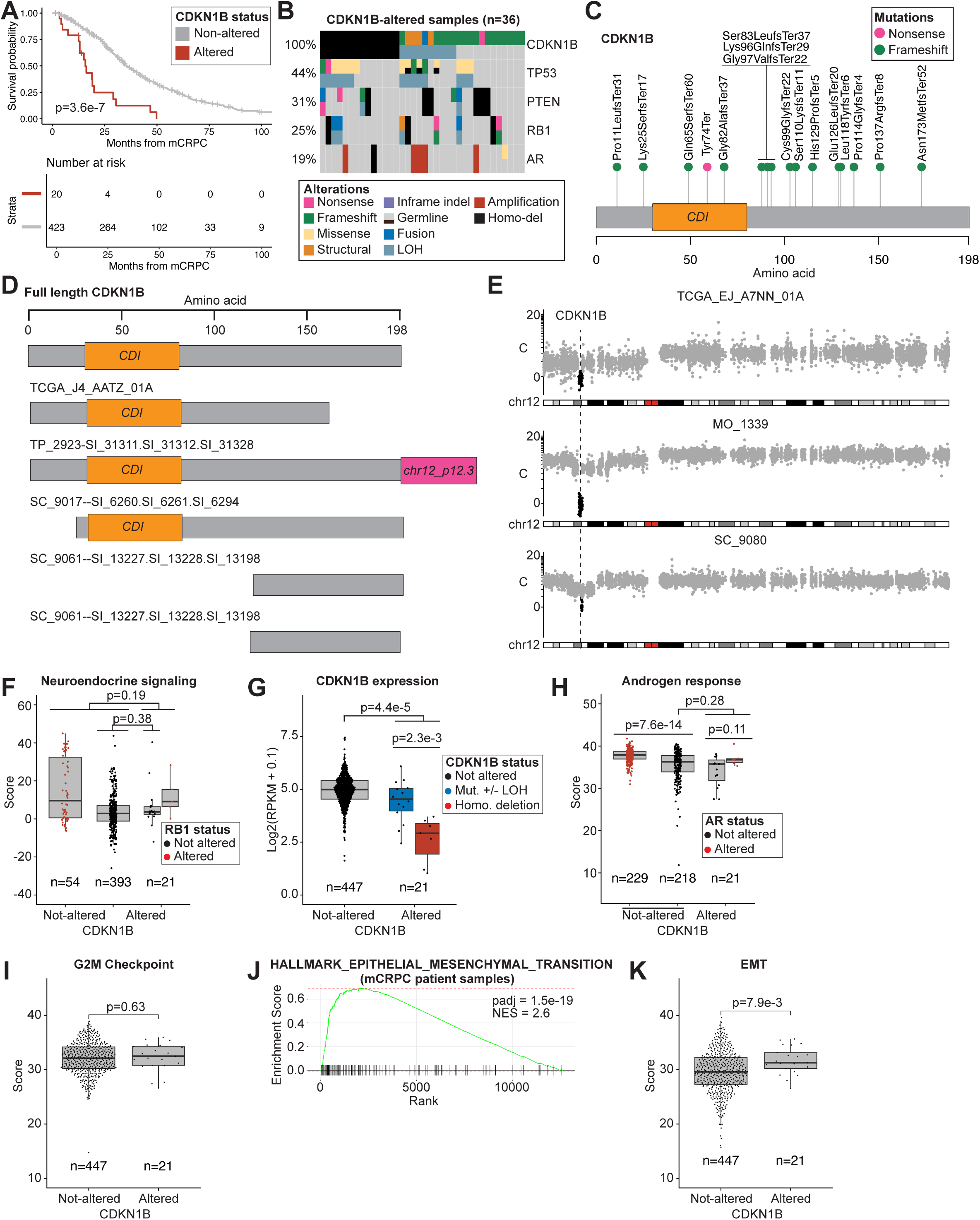
*CDKN1B* alterations drive an epithelial mesenchymal transition in lethal prostate cancer. **A.** Kaplan-Meier plot of overall survival from the time of metastatic castration-resistant prostate cancer for patients with CDKN1B alterations compared to patients without CDKN1B alterations. Significance determined using log-rank test. **B.** Oncoprint of 36 high-quality *CDKN1B*-altered samples. Concurrent alterations in *RB1, TP53, PTEN*, and *AR* shown. **C.** Amino acid locations of *CDKN1B* 16 mutations. 1 splice alteration (chr12:12718316_T/G; SC_9170) not shown. CDI=Cyclin-dependent kinase inhibitor domain. **D.** Structural alterations in *CDKN1B*. UTR=untranslated region. **E.** Copy number plot of chromosome 12 highlighting *CDKN1B* focal deletions. C=copy number. **F-I.** High-quality metastatic castration-resistant prostate cancer (mCRPC) samples stratified by CDKN1B alteration status with **F)** Neuroendocrine gene signature scores, **G)** CDKN1B gene expression, **H)** Androgen signaling gene signature scores, and **I)** G2M Checkpoint gene signature scores. Significance assessed using Wilcoxon rank sum test. Abbreviations: mutation (Mut.), loss of heterozygosity (LOH). **J.** Gene set enrichment analysis for epithelial-mesenchymal transition in *CDKN1B*-altered mCRPC samples. NES=normalized enrichment score. Adjusted p-value shown. **K.** Epithelial mesenchymal transition gene signature scores in high-quality mCRPC samples stratified by CDKN1B alteration status.

Notably, *CDKN1B* alterations were frequently monoallelic (12/36, 33%) (**Figure 4B**). This mirrors the heterozygous truncating mutations found in small-intestine neuroendocrine tumors ^73^ and hairy cell leukemia ^74^, two malignancies where *CDKN1B* is recurrently mutated. Stratification of *CDKN1B*-altered patients by allelic status (monoallelic vs. biallelic) or alteration class (mutation vs. homozygous deletion) showed no significant differences in OSmcrpc (p=0.72, p=0.82) (**Figure S4A-B**). We next examined how *CDKN1B* alterations overlap with other common alterations in mCRPC, finding that they were significantly enriched for *SPOP* alterations (generally associated with favorable outcomes) (p=0.015) but depleted of *AR* alterations (p=0.039) and ETS fusions (p=0.034) (**Figure S4C**). Frequent co-occurring alterations included *TP53* (44%), *PTEN* (31%), and *RB1* (25%) (**Figure 4B**).

Given the established role of *CDKN1B* as a driver in neuroendocrine malignancies of the small intestine and pancreas ^75^, we asked whether these alterations promote neuroendocrine differentiation in mCRPC. As previously reported ^22^, global genomic profiles were comparable between adenocarcinoma and non-adenocarcinoma subtypes (**Figure S4D**), with the exception of *RB1* alterations which were strongly associated with neuroendocrine histology (65% vs 11%, p=4.8e-11) (**Figure S4E**). In contrast, *CDKN1B* alterations showed no enrichment in the neuroendocrine subtype (4% vs 10%, p=0.50) (**Figure S4F**). Notably, of the 36 samples with *CDKN1B* alterations, only 2 (6%) had neuroendocrine differentiation at the time of sample collection (**Table S8**). Additional clinical review revealed evidence of neuroendocrine differentiation later during follow-up in 4 cases (**Table S8**). We observed no association between *CDKN1B* status and neuroendocrine gene expression in mCRPC (p=0.19), and this remained non-significant after excluding *RB1*-altered tumors (p=0.38) (**Figure 4F**). Overall, only 17% (6/36) cases developed neuroendocrine features, almost always in the presence of concurrent *RB1* (5/6) or *TP53* (4/6) alterations.

*CDKN1B* alterations significantly reduced *CDKN1B* expression in mCRPC (p=4.4e-5), with homozygous deletions driving deeper transcriptional repression than mutations (p=2.3e-3) (**Figure 4G**). We observed a robust, dosage-dependent correlation between *CDKN1B* copy number and expression across both localized (R=0.37) and castration-resistant (R=0.45) disease, indicating that genomic loss results in functional protein deficits. This mirrors the biology of genetically engineered mouse models, in which prostate tumorigenicity is governed by *Cdkn1b* levels and heterozygous loss suffices to drive carcinogenesis ^76^.

To define the transcriptional consequences of *CDKN1B* loss, we compared gene expression between altered and non-altered tumors. Consistent with the prevalence of truncating events, *CDKN1B* itself emerged as the most significantly downregulated gene in both the localized (FDR=7.0e-3, logFC= -0.71) and castration-resistant (FDR=7.9e-5, logFC= -1.1) settings (**Figure S5A, Table S9**). Despite being depleted of *AR* genomic alterations (p=0.034), *CDKN1B*-altered tumors maintained AR signaling levels comparable to *AR* non-altered tumors (p=0.28) (**Figure 4H**). Importantly, while we observed no significant change in proliferative signals (p=0.63) (**Figure 4I**), *CDKN1B*-altered tumors displayed a marked upregulation of the epithelial mesenchymal transition (EMT) (padj=1.5e-19, NES= 2.6) (**Figure 4J**, **Table S10**). In *CDKN1B*-altered mCRPC patient samples, *SHH*, a principal regulator of EMT, was the top upregulated gene (**Figure S4A**); consistent with this, these samples displayed significantly higher levels of EMT pathway activity (**Figure 4K**).

In mice biallelic *Cdkn1b* deletion alone is insufficient to drive prostate cancer formation, and requires concurrent heterozygous *Pten* inactivation ^77^, while mechanistically *Cdkn1b* loss abrogates *Akt1* transgene induced senescence ^78^. Notably, an analogous mechanism for *Cdkn1b* loss is proposed for *Shh*-driven medulloblastoma ^79^. Therefore, to further interrogate the effect of *CDKN1B* loss, we performed RNA-sequencing of *RB1* wild-type but *PTEN*-deficient LNCaP cells following siRNA-mediated *CDKN1B* knockdown (**Figure S5B**). This in vitro model faithfully recapitulated the unique clinical phenotype. *CDKN1B* loss alone was insufficient to induce neuroendocrine gene expression (**Figure S5C**), but instead maintained a robust AR transcriptional program (**Figure S5B,D**) without accelerating proliferation (**Figure S5E**). Crucially, the model confirmed the invasive potential observed in patients, showing a significant upregulation of EMT signaling (p=5.8e-3, NES=1.6) (**Figure S5F**).

Collectively, these data define *CDKN1B* loss as a uniquely lethal prostate cancer subtype, driving an AR-intact, mesenchymal phenotype biologically distinct from neuroendocrine transdifferentiation associated with *TP53*/*RB1* co-alteration.

### *TP53* and whole-genome duplication cooperate to drive lethal genomic instability

Across both Discovery and SU2C cohorts, the median OSmcrpc for *TP53*-altered patients was 28.7 months, significantly worse than the 44.5 months observed for non-altered patients (p=6.1e-9) (**Figure 5A**). While this reinforces the negative prognostic value of *TP53* alterations in metastatic castration-resistant prostate cancer, our understanding of the mechanisms by which *TP53* loss drives lethality in prostate cancer remains incomplete.

**Figure 5:**
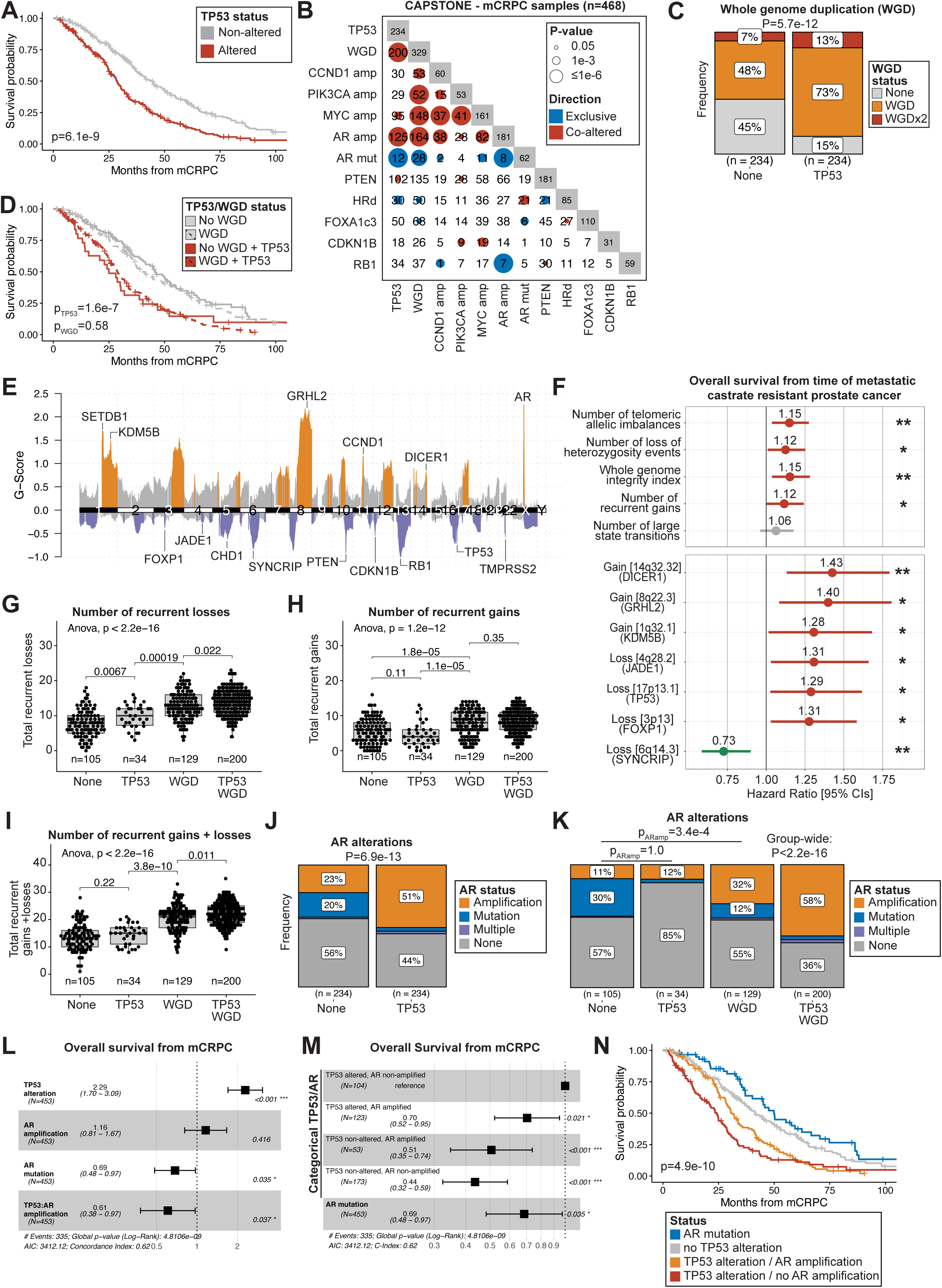
*TP53* loss facilitates recurrent copy number gains via whole-genome duplication in lethal prostate cancer. **A.** Overall survival from the time of mCRPC (OSmcrpc) by TP53 status. Significance determined using log-rank test. **B.** Co-occurrence or mutual exclusivity of notable genomic alterations in 468 metastatic castration-resistant prostate cancer (mCRPC) samples. Significance is represented by dot size. Abbreviations: Whole genome duplication (WGD), FOXA1 class III mutation (FOXA1c3). **C.** Whole genome duplication (WGD) status by *TP53* alteration status in mCRPC samples. Significance determined using chi-squared test. **D.** OSmcrpc by TP53 and WGD status. P-values represent significance from a multivariable cox proportional hazards model with TP53 and WGD for OSmcrpc. **E.** GISTIC plot for 1,160 prostate cancer samples. Recurrently gained/lost peaks shown in orange/purple respectively. Select genes within recurrently altered peaks highlighted. **F.** Hazard ratio for WGD, scaled measures of CIN, total number of recurrent gains, and select recurrently altered regions versus OSmcrpc in the pooled DIS/SU2C cohorts. Point estimates represent hazard ratios, lines represent 95% confidence intervals. Significance: * <0.05, ** <0.01, ***<0.001. **G-I.** The number of **G)** recurrent losses, **H)** recurrent gains, and **I)** recurrent gains and losses for mCRPC samples stratified by *TP53* alteration and WGD status. ANOVA and Wilcoxon tests used to determine significance. **J.** AR amplification and AR mutation status by *TP53* alteration status in mCRPC samples. Significance determined using chi-squared test. **K.** AR mutation and amplification status by *TP53* alteration status and WGD status. Chi-squared tests used to determine significance. **L-M.** Cox proportional hazard model incorporating, **L**) *TP53* alteration status, *AR* mutation status, *AR* amplification status, and the interaction between *TP53* and *AR* amplification, and **M**) AR mutation status and categorical representation of TP53 alteration status and AR amplification status. While (L) facilitates TP53 non-altered, AR non-amplified, AR non-mutated as the reference group, (M) facilitates TP53-altered, AR non-amplified, AR non-mutated tumors as the reference group for comparison. **N.** OSmcrpc between patients stratified into 4 groups: patients with an *AR* mutation irrespective of *TP53* alteration status, patients without a TP53 alteration, patients with a *TP53* alteration and an AR amplification, and patients with a *TP53* alteration but without an *AR* amplification. Significance determined using log-rank test.

First, we sought to separate the direct impact of *TP53* alterations from their role in maintaining genome integrity ^80^, and facilitating copy-number gains and losses. We observed that *TP53*-altered tumors were significantly enriched for whole-genome duplication (WGD) events (p=5.7e-12) (**Figure 5B**, **Methods**). One or more rounds of whole genome duplication were observed for 85% of *TP53*-altered compared to 55% non-altered tumors (**Figure 5C**). Although the presence of WGD itself was associated with poor OSmcrpc (p=0.022, univariate), having multiple WGD events did not impact prognosis further (p=0.62) (**Figure S6A**). Notably, the association between WGD and prognosis was driven primarily by the association of WGD with TP53 alterations rather than the presence of WGD alone (p=0.58, multi-variate) (**Figure 5D**).

To disentangle the direct downstream effects of *TP53* alterations from those mediated by WGD, we analyzed how the interaction between these variables influenced genomic integrity. We focused on global metrics of CIN (wGII, nLST, nTAI, nLOH) as well as the frequency of gains and losses at “driver” loci with recurrent copy-number alterations (**Figure 5E**, **Table S11**). Notably, measures of CIN were almost uniformly associated with worse OSmcrpc (**Figure 5F**), and consistently highest in *TP53*-altered whole genome duplicated tumors (**Figure S6B-E**). Stratifying patients by *TP53* and WGD status revealed that *TP53* alterations alone were sufficient for the accumulation of recurrent genomic losses (p=6.7e-3), but not recurrent gains (p=0.11) (**Figure 5G-H**). Rather, WGD was required for the accumulation of recurrent genomic gains (p=1.8e-5) (**Figure 5G**). However, both *TP53* alterations and WGD were required for tumors to tolerate the highest total burden of recurrent copy number alterations (**Figure 5I**).

We proceeded to dissect this cumulative burden, aiming to isolate specific alterations associated with patient outcome. Screening copy-number gains and losses at recurrently altered regions, we identified multiple prognostic loci (**Figure 5F**, **Figure S6F**). High-risk loci included losses at 17p13.1 (*TP53*), 4q28.2 (*JADE1*), and 3p13 (*FOXP1*), alongside gains at 14q32.32 (*DICER1*), 8q22.3 (*GRHL2*), and 1q32.1 (*KDM5B*). Conversely, loss of 6q14.3 (*SYNCRIP)* ^81^ emerged as the sole recurrent alteration associated with improved prognosis (**Figure 5F**, **Figure S6F**).

Collectively, these data show that *TP53* alterations facilitate the evolution of lethal phenotypes with WGD acting as the proximal driver of genomic instability facilitating the accumulation of copy-number gains at risk-associated loci. *TP53* loss serves a dual role: it enables WGD and its amplification program while simultaneously permitting the accumulation of extensive genomic deletions that WGD alone cannot sustain.

### Distinct AR alterations drive favorable outcomes in *TP53*-altered and non-altered contexts

To investigate whether *TP53* alterations promote specific copy number changes within driver genes, we screened for co-occurrences and identified *AR* amplification as a dominant signal (**Figure 5B**). Strikingly, we found that tumors exhibited a dichotomy in the type of *AR* alteration depending on *TP53* status; *TP53*-altered tumors were significantly enriched for *AR* amplifications (53% vs 24%, p=1.1e-10), but depleted of activating *AR* mutations (5% vs 21%, p=4.5e-7) (**Figure 5J**). We identified WGD as the crucial mediator of this phenotype. Stratification revealed that WGD, rather than *TP53* status itself, was the primary determinant of *AR* focal amplification status (32% vs 11%, p=3.4e-4) (**Figure 5K**). Specifically, *TP53* alterations alone were not associated with increases in *AR* copy number (p=0.35), whereas WGD alone showed a significant association (mean: 10.5 vs 3.4, p=1.2e-12) (**Figure S6G**). Notably, the association between WGD and focal copy number gains above baseline ploidy of each chromosome extended to other key oncogenes, including CCND1 (19% vs 6%, p=6.2e-3), MYC (44% vs 9%, p=4.2e-9), and PIK3CA (19% vs 0%, p=6.2e-3) (**Figure S6H-J**).

Since our prior analysis has shown that canonical AR signaling is associated with favorable outcomes (**Figure 3C**) we proceeded to dissect the joint impact of *TP53* status and *AR* alterations on patient survival. We determined that androgen signaling was elevated in tumors with *AR* alterations, independent of the alteration type (**Figure S6K**). Additionally, we found that androgen signaling was associated with improved survival in *TP53*-altered tumors (HR: 0.66 [0.57-0.77], p=8.6e-8), but not TP53 non-altered tumors (HR: 0.89 [0.77-1.02], p=0.081) (**Figure S6L,M**). To determine whether genetic *AR* alterations were similarly prognostic we stratified patients by their *TP53* and *AR* alteration statuses (**Figure S6N**). This revealed a clear trend linking *AR* alterations to better outcomes in *TP53*-altered contexts (p=0.014).

To test this formally, we constructed a survival model incorporating *TP53* alterations, *AR* amplifications, and *AR* mutations as covariates, explicitly testing for an interaction between *TP53* status and *AR* amplification status. While *AR* mutations predicted longer survival independent of *TP53* status (HR: 0.69 [0.48-0.97], p=0.035), AR amplifications conferred an improved prognosis in *TP53*-altered tumors (HR: 0.70 [0.52-0.95], p=0.021) but not *TP53* non-altered tumors (HR: 1.16 [0.81-1.67], p=0.42) (**Figure 5L,M**). This may be partially explained by the depletion of *RB1* alterations in *AR*-amplified samples (p<1e-6) (**Figure 5B**). Rather than accumulating aggressive, neuroendocrine phenotypes these tumors are ‘locked-in’ to targetable, androgen-dependent signaling.

Taken together, these findings categorize patients into 4 prognostic groups: those with *TP53* alterations and without *AR* alterations have the worst outcomes, while those with AR mutations (typically in *TP53* wild-type tumors) have the best survival (**Figure 5N**). Patients with TP53 alterations and AR amplifications and those without TP53 alterations represent intermediate groups. For *TP53*-altered tumors, *AR* amplification likely identifies a subset of cancers that remain dependent on AR signaling rather than undergoing lineage plasticity. In contrast, *AR* mutations are more frequently detected in *TP53* non-altered tumors and are associated with the most favorable patient survival. Thus, distinct genomic mechanisms—AR amplification in unstable tumors and hotspot mutations in stable ones—converge to mark an AR-dependent cell state with improved outcomes in lethal disease.

### Development and validation of a unified multi-omic classifier for robust risk stratification in lethal prostate cancer

Building on our characterization of independent genomic and transcriptomic prognostic features, we sought to synthesize these insights into a unified multi-omic classifier for risk stratification. We employed a robust statistical workflow to identify a gene set associated with OS in mCRPC. As neuroendocrine transdifferentiation is known to be strongly associated with poor prognosis in mCRPC ^19,42,43,82,83^, we removed 15 samples with molecular evidence of neuroendocrine biology to isolate prognostic signals distinct from this dominant and well-understood aggressive phenotype (**Figure 6A**). Additionally, to better account for sequencing platform batch effects, the SU2C cohort was sub-stratified into 142 patients profiled with capture RNA sequencing ^84^ (SU2C-capture) and 101 patients profiled with Poly(A) RNA sequencing (SU2C-polyA). In total, 175 patients were available for model development, and 243 patients were available for subsequent independent validation.

**Figure 6:**
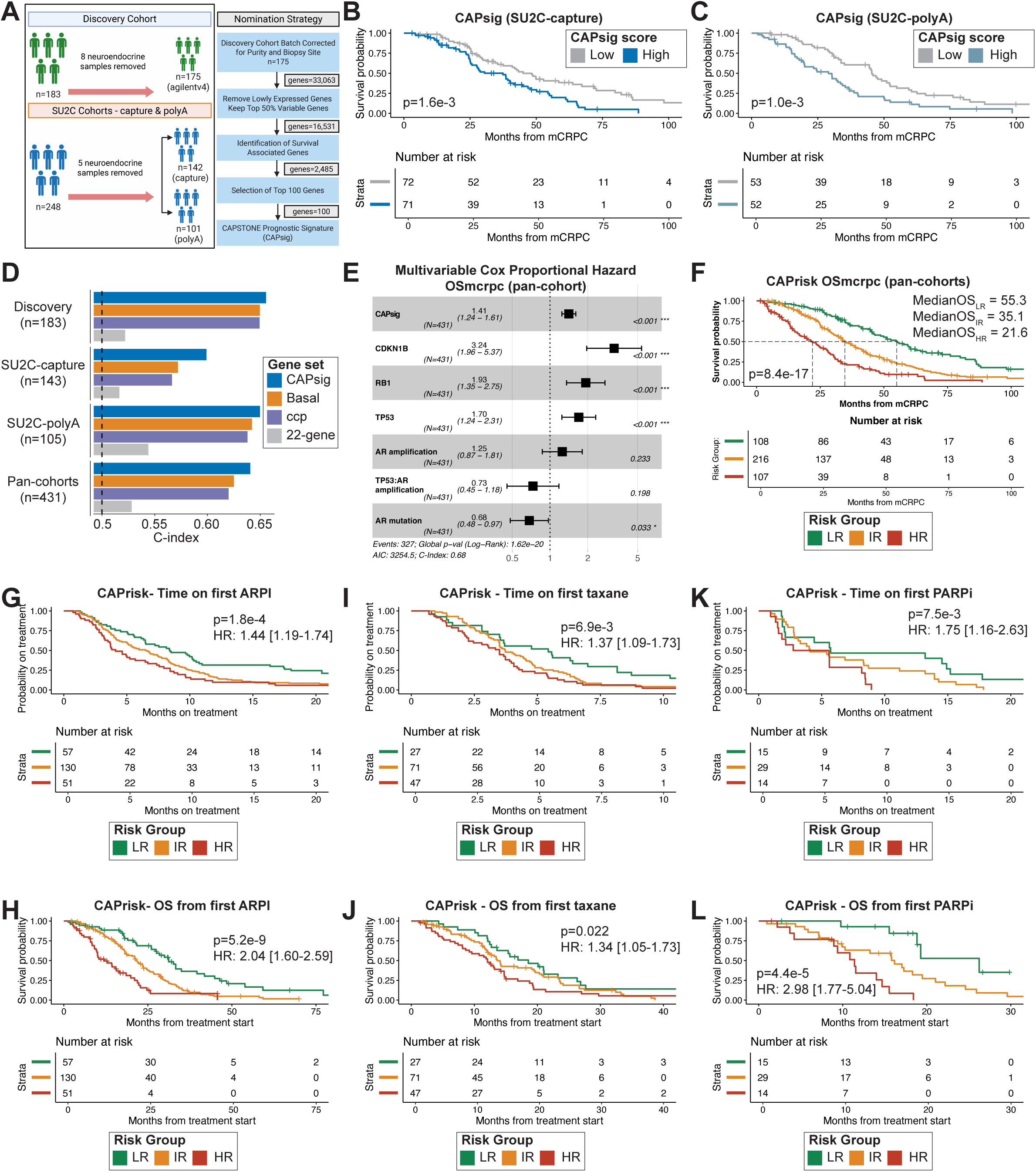
Development and application of a multi-omic classifier in lethal prostate cancer. **A.** Overview of the methodology used to generate a prognostic gene set in non-neuroendocrine metastatic castration-resistant prostate cancer (mCRPC). **B-C.** CAPsig performance for overall survival from the time of metastatic castration-resistant prostate cancer (OSmcrpc) in the SU2C capture B) and poly A) subsets with neuroendocrine samples included. Significance determined using log-rank tests. **D.** Performance (C-index) comparison between the prognostic gene set and other gene sets from external prognostic classifiers in prostate cancer. Performance stratified pan-cohort and by constituent cohorts. **E.** Multivariable Cox Proportional Hazards model for OSmcrpc and CAPsig and *CDKN1B*, *TP53*, AR, and *RB1* alteration status pan-cohort. **F.** Kaplan-Meier plot for CAPrisk groups and OSmcrpc. Significance assessed via log rank test. **G-H**. Kaplan-Meier plot for CAPrisk groups and Androgen Receptor Pathway Inhibitors (ARPIs) for **G**) time on treatment (ToT), and **H**) overall survival from treatment start (OStxt). **I-J**. Kaplan-Meier plot for CAPrisk groups and taxane chemotherapy for **I**) ToT, and **J**) OStxt. **K-L**. Kaplan-Meier plot for CAPrisk groups and PARP inhibitors (PARPi) for **K**) ToT and **L**) OStxt. For all kaplan-meier plots: LR=low-risk, IR=intermediate-risk, HR=high-risk, significance assessed using cox proportional hazards model unless otherwise noted.

To develop the prognostic gene set, we analyzed 16,531 candidate genes (**Figure 6A**), utilizing dimensionality reduction to identify two distinct prognostic groups (**Methods**). Through optimization of the differentially expressed features between these states, we isolated a final 100-gene set termed the CAPSTONE prognostic signature (CAPsig) (**Table S12**). CAPsig was enriched for biological processes related to proliferation such as mitotic nuclear division (GO:0140014; p-adj = 3.3e-7) (**Figure S7A**), but comprised a range of molecular functions, notably tissue remodeling (*MMP1*, *MMP7*), basal luminal markers (*KRT5*, *KRT15*), mitotic spindle assembly (*KIF18A*, *KIF2C*), pro-inflammatory cytokines (*CXCL8*, *CXCL17*), and histones (**Table S12**).

High expression of CAPsig was associated with poor OSmcrpc across both SU2C-capture (p=1.6e-3), SU2C-polyA (p=3.5e-3), and the Discovery cohorts (p=3.5e-7) (**Figure 6B-C**, **S7B**). We benchmarked CAPsig against constituent genes from three of the most robustly validated prognostic classifiers developed for hormone-sensitive prostate cancer: the cell cycle progression (CCP) signature ^85^, the basal signature ^24–26^, and the 22-gene prognostic signature ^32^, as well as prognostic signatures from (**Figure 2**) (**Methods**). This direct comparison revealed that CAPsig yielded superior prognostic power in each of the three individual cohorts, as well as in the pooled cohort (**Figure 6D**, **Figure S7C**). Notably, while it correlated with established basal and CCP signatures (**Figure S7D**), CAPsig represented a distinct biological profile, sharing only 16 constituent genes with these classifiers (**Figure S7E**).

Motivated by these findings we sought to combine CAPsig with prognostic genomic features into a composite multi-omic risk classifier. First, to determine if CAPsig captures biological risk distinct from established genomic drivers, we evaluated its prognostic utility alongside alterations in *TP53*, *RB1*, *AR,* and *CDKN1B*. Using multivariable analysis, we found that CAPsig was associated with OSmcrpc independent of *CDKN1B*, *RB1*, *AR,* and *TP53* alteration status (HR: 1.41 [1.24-1.61], p=2.8e-4) (**Figure 6E**). Notably, this remained significant when stratified into the Discovery (HR: 1.33 [1.11-1.59], p=1.6e-3), SU2C-capture (HR: 1.43 [1.08-1.87], p=0.011, and SU2C-polyA (HR: 1.54 [1.16-2.03], p=2.7e-3) cohorts.

Next, we combined CAPsig and the prognostic alterations into a multi-omic risk classifier (CAPrisk). Stratification into three distinct risk groups via leave-one-out cross-validation i.e. providing cross-validated within-cohort estimates (**Methods**) revealed significantly divergent outcomes (p=8.4e-17) (**Figure 6F**). Median OSmcrpc for CAPrisk low-risk (LR) patients was 55.3 months compared to 35.1 months for CAPrisk intermediate risk (IR) and 21.6 months for CAPrisk high-risk (HR) patients.

CAPrisk groups remained significant when stratified into the Discovery (p=1.2e-11), SU2C-capture (p=1.5e-3), and SU2C-polyA (p=7.3e-4) cohorts.

HR patients were enriched for *CDKN1B* alterations (p=4.1e-11), *RB1* alterations (p<2.2e-16), and *TP53* alterations (p<2.2e-16) (**Table S13**). We also observed that HR patients were frequently metastatic at diagnosis (53/107, 50%) and had higher rates of ISUP grade 5 disease at time of diagnosis than LR patients (47% vs 24%, p=0.012) (**Table S13**). Additionally, rates of neuroendocrine histology were greater in HR patients (10/107, 9%) compared to IR (1/216, 0%) and LR patients (0/108, 0%) (p=1.8e-6) (**Table S13**).

### CAPrisk stratifies clinical outcomes across standard-of-care therapies

To evaluate CAPrisk as a potential instrument of precision medicine, we investigated its utility in stratifying response to standard-of-care therapies. We assigned risk groups to tumor samples collected prior to or within 12 months from therapy exposure (**Methods**) and evaluated Time on Therapy (ToT) and treatment-specific Overall Survival (OStxt) across major therapeutic classes including ARPIs, Taxanes, PARPi, and Radium-223. CAPrisk demonstrated robust stratification across distinct therapeutic classes.

Within patients receiving first-line ARPIs (n=238), we found that an increase in risk group was associated with a significantly shorter ToT (HR: 1.44 [1.19-1.74], p=1.8e-4) and OStxt (HR: 2.04 [1.60-2.59], p=5.2e-9) (**Figure 6G-H**). LR patients were enriched for *AR* mutations (39% vs 5%, p=3.5e-9), whereas HR patients were enriched for *AR* amplifications (31% vs 9%, p=1.5e-4) (**Table S13**). Interestingly, despite the high rates of AR alterations in the HR group, canonical androgen signaling was significantly reduced relative to both IR and LR patients (**Figure S7F**), which remained significant after removal of neuroendocrine samples (**Figure S7G**). This suggests that HR tumors either activate non-canonical AR signaling or exhibit heightened lineage plasticity, facilitating therapy-resistant disease progression.

In patients receiving first-line taxane chemotherapy (n=145), increasing risk was associated with shorter ToT (HR: 1.37 [1.09-1.73], p=6.9e-3) and OStxt (HR: 1.34 [1.05-1.73], p=0.022) (**Figure 6I-J**). These associations were even more pronounced for patients receiving first-line PARPi (n=58) for both ToT (HR: 1.75 [1.16-2.63], p=7.5e-3) and OStxt (HR: 2.98 [1.77-5.04], p=4.4e-5) (**Figure 6K-L**). Notably, the benefit observed in LR patients receiving PARPi held despite rates of HRd comparable to HR patients (p=0.27) (**Table S13**). Similarly, rates of *BRCA1/2* and *PALB2* alterations were comparable between LR and HR groups (**Table S13**). In light of approvals for PARPi use irrespective of HRd status ^86^, CAPrisk may prove useful in further substratifying homologous recombination proficient patients benefiting from PARPi. Additionally, although limited by sample size, we also observed a significant relationship between risk groups and both ToT and OStxt for Radium-223 (n=35, p=1.5e-3, 1.7e-3) (**Figure S7H-I**).

Overall, CAPrisk delineates a survival difference of nearly 3 years between high- and low-risk groups, stratifying outcomes across all major standard-of-care therapies.

### Therapeutic implications of recurrent genomic alterations in lethal prostate cancer

In contrast to the wealth of genomic datasets, there is a relative scarcity of hypothesis-generating studies pairing genomic alterations with treatment-specific outcomes in lethal prostate cancer. We leveraged information for 26 recurrent genomic alterations with treatment information across 7 therapeutic regiments to uncover novel associations with standard-of-care treatments (**Figure 7A**).

**Figure 7:**
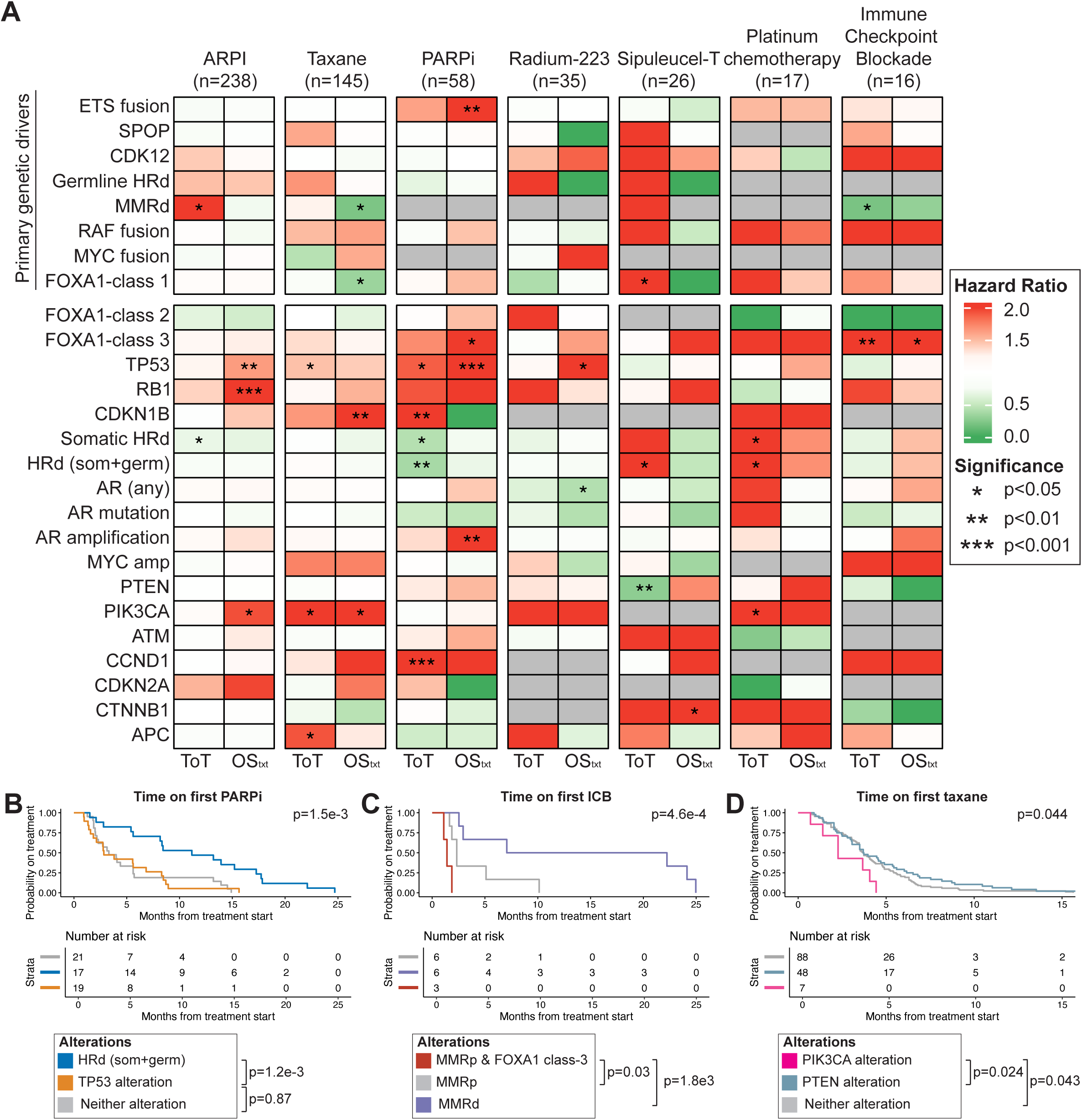
Therapeutic implications of recurrent genomic alterations in lethal prostate cancer. **A.** Heatmap of select recurrent genomic alterations and association with time on therapy (ToT) and overall survival from time of treatment start (OStxt) for therapies including Androgen Receptor Pathway Inhibitors (ARPIs), taxane chemotherapy, PARP-inhibitors (PARPi), Sipuleucel-t, platinum chemotherapy, and immune checkpoint blockade (ICB). **B-D.** Kaplan-Meier plot of **B)** ToT with PARPi stratified by homologous recombination deficiency status and *TP53* status, **C)** ToT with ICB stratified by mismatch repair (MMR) status and *FOXA1* class III alteration status, and **D)** ToT with taxane chemotherapy stratified by *PIK3CA* and *PTEN* alteration status. Significance assessed using log-rank tests.

Consistent with their limited prognostic value in mCRPC (**Figure 2**), primary genetic drivers showed relatively few significant associations with time on therapy (**Figure 7A**). The exceptions included known responses to immune checkpoint blockade (ICB) and PARPi for MMRd and HRd tumors, respectively ^86,87^. In this cohort MMRd prostate cancer was associated with prolonged ToT for ICB (HR^ToT^: 0.19 [0.051-0.73], p=0.016). Similarly, we found a trend between germline (gHRd) and improved ToT for PARPi (HR^ToT^: 0.66 [0.31-1.39], p=0.27). This association reached significance for total HRd (somatic and germline) (HR^ToT^: 0.37 [0.21-0.66], p=1.1e-3).

We next examined other alterations acquired during prostate cancer progression and systemic treatment. We confirmed that *RB1* alterations associated with shorter OStxt on ARPIs (HR^OStxt^: 2.15 [1.36-3.39], p=1.0e-3) ^19^. Further, we observed strong associations between *TP53* alterations and poor outcomes following ARPIs (HR^ToT^: 1.63 [1.22-2.21], p=1.2e-3) ^88,89^, taxane chemotherapy (HR^ToT^: 1.47 [1.05-2.07], p=0.025), Radium-223 (HR^OStxt^: 2.35 [1.07-5.17], p=0.034), and PARPi (HR^ToT^: 1.89 [1.07-3.36], p=0.030; HR^OStxt^: 3.25 [1.68-6.31], p=5.0e-4). For PARPi specifically, this negative prognostic signal was driven primarily by the mutual exclusivity between *TP53* alterations and HRd (**Figure 7B**).

Although our sample size was constrained by the analysis of rare alterations, we uncovered several associations warranting future investigation. We found that *CDKN1B* alterations conferred poor outcomes following PARPi (HR^ToT^: 28.0 [2.54-308], p=6.5e-3), and taxane chemotherapy (HR^OStxt^: 2.71 [1.30-5.62], p=7.6e-3). Additionally, we identified a robust link between *FOXA1* class-3 alterations (gains) and poor response to ICB (HR^ToT^: 15.6 [2.42-99.7], p=3.8e-3; HR^OStxt^: 9.58 [1.57-58.5], p=0.014). Patients harboring *FOXA1* class-3 alterations exhibited significantly worse ToT and OStxt compared to both MMRd and MMR-proficient controls (**Figure 7A**, **7C**). Lastly, *PIK3CA* alterations associated with poor outcomes on first-line taxane chemotherapy (HR^ToT^: 2.33 [1.07-5.04], p=0.032; HR^OStxt^: 2.69 [1.17-6.22], p=0.020) while *PTEN* alterations showed no association with taxane resistance, suggesting *PIK3CA* as the specific driver of this phenotype (**Figure 7D**).

Overall, by identifying candidate biomarkers of poor therapeutic outcomes, this analysis bridges genomic profiling to clinical utility and offers hypotheses for designing future biomarker-driven trials.

## DISCUSSION

In this study, we established CAPSTONE, a multi-institutional clinicogenomic atlas of 1,331 prostate cancer samples, and applied a rigorous discovery and validation framework to systematically define the genomic determinants of lethal disease. We distinguish the specific roles of drivers across disease progression: notably identifying *MYC* gene fusions as truncal events that drive rapid progression to metastatic disease akin to aggressive subtypes like *CDK12* loss, MMRd, and HRd. We also characterize *CDKN1B* alterations as a distinct driver of poor survival in the metastatic castration-resistant (mCRPC) setting, matching the adverse prognostic impact of established markers *TP53* and *RB1*. To translate these diverse molecular insights into a unified framework for clinical risk stratification, we created CAPrisk, a multi-omic prognostic classifier that integrates a novel transcriptomic risk signature (CAPsig) with genomic alterations to accurately stratify patient outcomes. Finally, by contextualizing these risk profiles with longitudinal clinical data, we uncovered significant associations between specific genomic features and therapeutic responses, providing a rationale for precision oncology strategies in lethal prostate cancer.

Despite being detected in over 75% of tumors ^10^, and ongoing clinical trials ^90^, primary genetic drivers outside of HRd are not currently used for prognostication or to direct prostate cancer treatment. We show that select primary genetic drivers are linked to poor outcomes. Consistent with previous findings, *CDK12* alterations were enriched in patients who developed mCRPC ^13^ and associated with shorter time to the development of mCRPC ^67^. We also found that in patients who develop lethal disease, both *SPOP* mutations and ETS fusions are associated with both longer time to castration-resistance and longer overall survival from time of diagnosis. This is consistent with recent reports linking *SPOP* mutations with improved ADT sensitivity in the *de novo* mCSPC setting^64^. Importantly, providing a human genetic rationale for the efficacy of *MYC*-overexpressing mouse models ^53,54^, we distinguish *MYC* fusions from canonical amplifications as a unique class of primary genetic drivers.

The fusion of *MYC* to androgen-upregulated genes (*e.g.*, *ACPP*, *SLC45A3*, *NKX3-1*) reveal a novel mechanism by which tumor cells bypass AR-mediated repression of endogenous MYC gene expression, which normally restrains proliferation in both healthy prostate epithelium ^62^ and prostate cancer cells ^63^. By juxtaposing the *MYC* coding sequence downstream of AR-responsive promoters or enhancers, these fusions place *MYC* under direct androgenic control and convert a growth-suppressive output of canonical AR signaling into an oncogenic driver of prostatic transformation.

Taken together, these data show that while primary genetic drivers do not dictate survival outcomes in late-stage mCRPC, they are fundamental drivers of the initial disease trajectory. Their impact is predominantly concentrated in the hormone-sensitive setting, where they serve as vital biomarkers for identifying patients at risk of accelerated progression versus those with indolent biology.

The cyclin-dependent kinase inhibitor *CDKN1B* (p27) has long been recognized as a critical gatekeeper of the G1/S checkpoint ^91^. Seminal immunohistochemical studies in the late 1990s first established that reduced p27 protein levels correlated with high Gleason grade and poor prognosis in localized disease ^92–96^. Subsequent murine modeling revealed that while *Cdkn1b* loss alone is insufficient to initiate prostate tumorigenesis, it synergizes with *Pten* deficiency to drive carcinoma development ^77^, and *Shh* in medulloblastoma ^79^. In humans, this tumor suppressive capability is underscored by germline variants linked to both familial ^97,98^ and sporadic prostate cancer risk ^99^.

Consistent with its non-canonical roles in regulating cell polarity ^78^ and established haploinsufficiency across multiple cancer types ^100–103^, we find that *CDKN1B* functions as a haploinsufficient somatic driver in mCRPC, where both monoallelic and biallelic alterations result in dosage-dependent expression loss and equally poor outcomes. Crucially, distinct from the neuroendocrine differentiation driven by *RB1* loss, we demonstrate through functional experiments that *CDKN1B* inactivation facilitates an AR-dependent metastatic progression by driving epithelial-mesenchymal transition. Our results thus bridge decades of genetic biological inquiry to cement *CDKN1B* alterations as a distinct and potent driver of poor survival in mCRPC.

To dissect the downstream genomic events by which *TP53* loss drives lethality, we establish a sequential model of genomic evolution wherein *TP53* loss catalyzes chromosomal instability and the selection of copy-number alterations at prognostic loci, including well-established oncogenes (*e.g. CCND1*) and tumor-suppressors (*e.g. FOXP1*). We further demonstrate that *TP53* status dictates the specific genomic mechanism of *AR*-driven castration resistance. *AR* hotspot mutations are enriched in *TP53* non-altered tumors, whereas *AR* amplifications are enriched in the *TP53*-altered setting.

Strikingly, both *AR* amplifications in the TP53-altered context and *AR* mutations in all contexts predict favorable survival. Thus, *AR* alterations identify tumors that remain dependent on androgen signaling, distinguishing them from the lethal *TP53*-altered/*AR* non-altered genotype, where the loss of lineage constraints permits aggressive plasticity.

To translate these diverse molecular insights into a clinical tool, we developed CAPsig, a novel transcriptomic signature that outperforms other prognostic classifiers in the lethal setting. By integrating CAPsig with the independent genomic prognostic drivers *TP53*, *RB1*, and *CDKN1B*, we established CAPrisk, a unified multi-omic framework that accurately stratifies patients into distinct risk groups with widely divergent survival outcomes (median OS: 55.3 vs 21.6 months). Crucially, this stratification goes beyond predicting survival to associating genetic alterations with treatment responses. It successfully identifies high-risk patients who are broadly resistant to standard therapies, including ARPIs, taxanes, and PARP inhibitors. Furthermore, building upon previous analyses which have focused on linking select genes or pathways (*TP53, AR, RB1*, PI3K, HRd) with time on treatment with ARPIs ^19,45,67,104,105^, our signal-finding analysis revealed distinct associations between genomic alterations and therapeutic outcomes, specifically highlighting the broad resistance conferred by *TP53* alterations, the resistance to taxanes linked to *CDKN1B* loss, and the lack of response to immunotherapy in *FOXA1* class-3 altered tumors.

Our study has limitations inherent to its retrospective, multi-institutional design. Patients were accrued over an extended period with Discovery patients having a median year of diagnosis of 2012. This introduces clinical heterogeneity, especially given the interval changes to standard of care treatments for metastatic prostate cancer. Further, despite the large cohort size, statistical power remains limited for rare genomic drivers. Furthermore, our reliance on single-site biopsies may underestimate molecular heterogeneity within a patient ^69^. We were also unable to directly compare our findings with existing prognostic models due to a lack of uniform laboratory data ^106^. Regarding primary drivers, the limited follow-up in the TCGA dataset presents a constraint. We assumed low-grade localized disease rarely progresses to mCRPC, but this assumption warrants confirmation.

Ultimately, prospective validation with unified enrollment criteria is essential. This step is critical to confirm the utility of these biomarkers and their associations with standard-of-care therapeutic outcomes.

## ACKNOWLEDGEMENTS

We would like to acknowledge Munna Hazime, Tanya Hammoud, and Pin-en Chiu for their work in the formation of the clinical database. A.M.C. is a Howard Hughes Medical Institute Investigator, A. Alfred Taubman Scholar, and American Cancer Society Professor. This work was supported by the National Cancer Institute Specialized Programs of Research Excellence grant (P50CA186786, A.M.C.).

## AUTHOR CONTRIBUTIONS

Conceptualization, R.J.R., L.H., W.C.J., J.J.A., Z.R., A.M.C., R.D., and M.C.

Methodology, R.J.R., M.G., N.H., M.S., R.D., M.C.

Investigation, R.J.R., M.G., A.P., and M.C.

Writing – Original Draft, R.J.R., A.P., W.C.J., J.J.A., Z.R., D.R., R.D., and M.C.

Writing – Review & Editing, R.J.R., A.P., M.S., A.K., M.H.H., T.C.W., W.C.J., J.J.A., Z.R., A.M.C., R.D., and M.C.

Funding Acquisition, W.C.J., J.J.A., Z.R., D.R., A.M.C., R.D., and M.C.

Resources, L.H., A.P., Y.W., M.M., X.C., P.P., M.C., D.S., S.Y., M.H.H., D.S., W.C.J., J.J.A., Z.R., D.R., A.M.C., R.D., and M.C.

Data Generation, R.J.R., L.H., A.P., Y.W., A.K., Z.R., and M.C.

Supervision, A.P., M.S., M.H.H., T.C.W., D.S., W.C.J., J.J.A., Z.R., D.R., A.M.C., R.D., and M.C.

## STAR METHODS

### Resource availability

#### Lead contact

Further information and requests for resources and reagents should be directed to and will be fulfilled by the lead contact, Marcin Cieslik.

#### Materials availability

All sequencing data generated as part of this study is deposited at dbGaP and publicly available as identified below. Any additional information required to reanalyze the data reported in this paper is available from the lead contact upon request.

#### Data and code availability

● Bulk DNA and RNA sequencing data have been deposited at dbGaP and are publicly available as of the date of publication.
● All original code has been deposited at Github (https://github.com/mctp/tpo-public-v2.5) and is publicly available as of the date of publication. DOIs are listed in the key resources table.
● Any additional information required to reanalyze the data reported in this paper is available from the lead contact upon request.

#### Experimental model and study participant details

##### Human participants

This study was approved by the Institutional Review Board of the University of Michigan (Michigan Oncology Sequencing Protocol, MI-ONCOSEQ, IRB HUM00067928, HUM00046018). Patients 18 years or older with a suspected diagnosis of prostate cancer were eligible for the study.

#### Method details

##### Integrative clinical sequencing

Histologic sections were evaluated for tumor content prior to sequencing. Nucleic acid preparation and integrative clinical sequencing (comprising exome sequencing and capture RNA-seq) was performed using standard protocols in our sequencing laboratory, which adheres to the Clinical Laboratory Improvement Amendments ^29^. In brief, tumor genomic DNA and total RNA were purified from the same sample using the AllPrep DNA/RNA/miRNA kit (Qiagen Cat#80224). Matched normal genomic DNA from blood, buccal swab or saliva was isolated using the DNeasy Blood & Tissue Kit (Qiagen Cat#69504). Targeted Exome-capture libraries of matched pairs of tumor and normal DNA were prepared as previously described ^29^. Transcriptome libraries were prepared from total RNA captured by human all-exon (Agilent SureSelect Human All Exon v4) probes. All samples were sequenced on an Illumina HiSeq 2000, HiSeq 2500, or NovaSeq 6000 (Illumina) in paired-end mode of reads at least 125 bp in length. The primary base call files were converted into FASTQ sequence files using the bcl2fastq converter tool bcl2fastq-1.8.4 or later.

#### Comprehensive genomic analysis pipeline

All genomic data processing and analysis has been performed using the TPO workflow (https://github.com/mctp/tpo-public-v2.5), which implements standardized pipelines for the analysis of DNA and RNA sequencing data, broadly following community best practices ^107^. Relevant algorithmic details are explained below, a general overview of its functions has been provided in prior publications ^69,108^.

#### Somatic variant calling

Total number of reads, percent of duplicated reads, and percent of chimeric reads were calculated for exome capture data using the same criteria as Picard ^109^. Other quality measures such as GC-bias ^110^ and read mappability were factored directly in the CNV and variant calling processes to minimize the effects of data quality on sensitivity of the calls. BBMap’s bbduk ^111^ was used to perform trimming of DNA sequencing paired-end reads. Next, the data was aligned to the GRCh38 reference using BWA-mem ^112^. Sentieon sort tool ^113^ was used to sort the reads in the BAM files. For somatic variants, all tumor samples were matched with normal tissue. TNscope ^113^, an improved somatic caller based on GATK Mutect2 ^114^ was used to call the somatic variants and calculate variant allele frequency (VAF) according to its probabilistic model, taking realignment and mapping ambiguity into account. The following settings were used in TNScope:

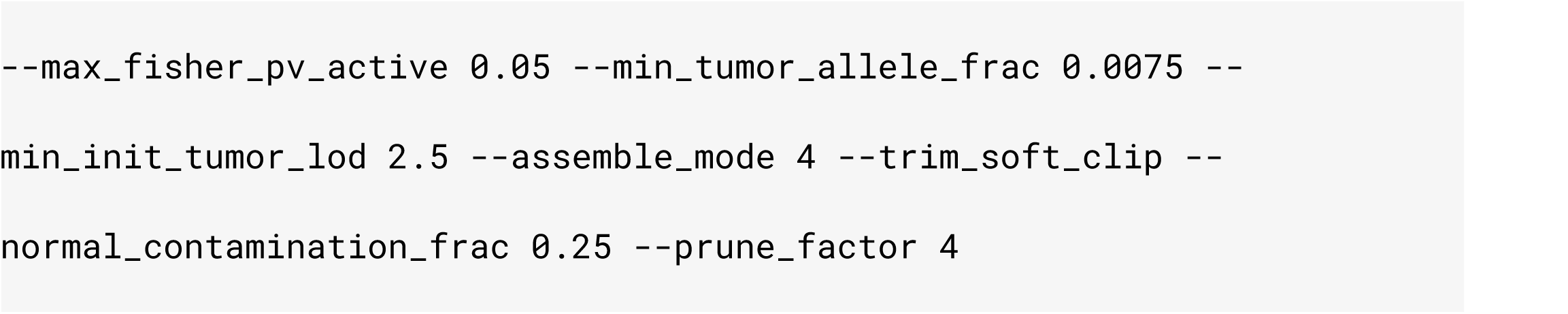

The called variants were annotated using VEP ^115^ and vcfAnno ^116^. All detected variants were further stringently filtered based on sequencing evidence (variant allele frequency, coverage, mutation likelihood TLOD and NLOD, strand bias, allele depth, multi-allelic variants), overlap in problematic regions including regions with low-mappability and repetitive sequence, and homopolymer repeats, and in ambiguous cases manually reviewed.

#### Copy number estimation

Copy-number analysis was performed by using whole-exome sequencing (WES) coverage data and variant calls based on the tumor DNA. CNVEX (https://github.com/mctp/cnvex) was used to estimate CNVs ^108,117^. Briefly, CNVEX estimates coverage within fixed genomic intervals and variant calls to compute B-allele frequencies (BAFs) at variant positions. Coverage values are then normalized for GC-bias using LOESS smoothing across targeted regions within the GC range of 0.3 and 0.7, using the span=0.5. All the GC normalized coverages and BAFs are then jointly segmented using a custom algorithm based on Circular Binary Segmentation (CBS) ^118^. The resulting segmented copy-number profiles were then subject to joint inference of tumor purity and ploidy and absolute copy number states, implemented in CNVEX, which is most similar to the mathematical formalism of ABSOLUTE and PureCN ^110,119^. Because the copy-number inference problem can have multiple equally likely solutions, further biological insights are necessary to choose the most parsimonious result. The solutions were reviewed by two independent field experts and the most likely solution was selected.

#### RNA-sequencing expression quantification

For RNA-seq data analysis, including read-trimming, alignment, post-processing, and quantification, we used the same methods as previously described ^108^. Briefly, adapter trimmed paired-end 150bp short reads were aligned using STAR version 2.4.0j to the GRCh38 reference supplemented with human oncogenic viruses ^120^. To improve the sensitivity of detecting fusion calls, synthetic single end reads were generated using bbduk from the trimmed paired-end reads. These reads were also aligned using STAR with optimized settings. After alignment, capture RNA-seq data were quantified using Kallisto ^121^. The paired-end and single-end RNA-seq alignments were used subsequently as an input to CODAC, a component of TPO designed to call fusions, perform additional QC and realign reads using Minimap and GMAP ^122,123^.

#### RNA-sequencing assay normalization

A minority of mCRPC samples within the CAPSTONE Discovery (n=5) and SU2C (n=132) cohorts were assayed via polyA rather than Agilent V4 capture RNA sequencing. To account for the batch effects attributable to the polyA assay method, we adjusted the RPKM values for the mCPRC samples profiled using polyA using linear regression in all analyses excluding those related to CAPrisk generation. Briefly, we fit a linear model on the batch (0=capture, 1=polyA) across all profiled genes. We then extracted the beta coefficients for each gene from this model and adjusted the log(RPKM) values for the polyA samples by this beta value. This ensured that RPKM values for all samples profiled using agilentV4 values were not altered. Select genes were checked before and after adjustment to ensure quality. This approach is based on statistical techniques utilized by ComBat ^124^.

#### Detection of gene fusions from RNA-sequencing

Detection of gene fusions was performed using the CODAC algorithm as previously described ^108,117^. Briefly, CODAC implements detection of genic and intergenic gene fusions based on both split- and discordant-reads as detected through chimeric read alignment using STAR ^120^. To maximize sensitivity, STAR is run separately using optimized settings in single-end and paired-end mode for overlapping and non-overlapping read pairs, respectively. The resulting alignments are merged, resulting in candidate fusion junctions identified from the STAR alignments are evaluated based on alignment properties to identify false-positive calls, including incorrect mappings, reference errors/differences, and non-genetic sources, such as circRNAs. The resulting call-set is further filtered against a manually-curated database of recurrent artifacts. STAR settings:

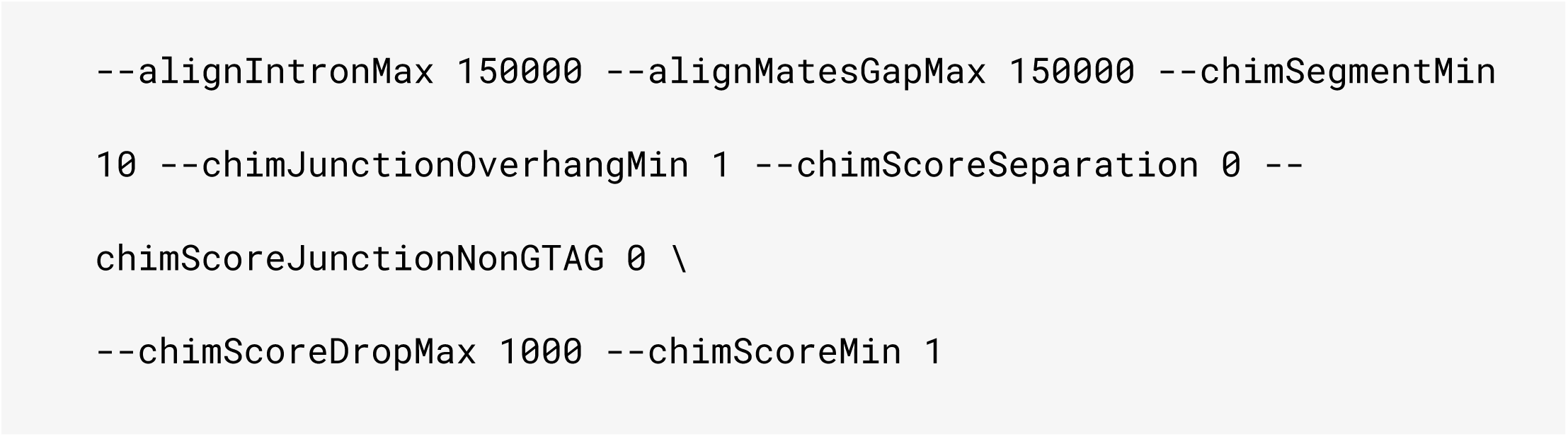

#### Annotation of focal copy number alterations

Gene-level homozygous deletions, loss of heterozygosity, and amplifications were computed using the following parameters:

● Amplifications (Copies ≥ 8 & Copies ≥ twice the basepair-weighted average copy number of the chromosome arm)
● Homozygous deletions: (Copies = 0 )
● Loss of heterozygosity: (Copies=1 | Minor allele copies =0 ) Where K is the number of minor alleles.

#### Identification of high-quality patient samples

Within CAPSTONE, we defined high-quality samples as those with matched sequencing from tumor RNA, tumor DNA, and non-cancerous tissue DNA. The paired DNA samples needed to be profiled via the same capture platform and after copy number analysis (see above) have sufficient sample coverage to allow fitting of a tumor purity/ploidy model.

#### Manual curation of alterations in important genes and pathways in prostate cancer

To determine whether an alteration was present in a gene, we used the available DNA and RNA-sequencing derived features including somatic mutations, gene fusions and structural alterations, genetic deletions / gains, and if applicable germline mutations. For select genes and pathways, alterations were manually reviewed across all samples to ensure annotation accuracy. These genes included:

● *ELK4, ERG, ETV1, ETV4, ETV5, FLI1, TMPRSS2 (*ETS gene fusion partners)
● *BRAF, RAF1* (RAF gene fusion partners)
● *RB1, CCND1, CDKN2A, CDKN1B* (cell cycle)
● *PIK3CA, PTEN* (PI3K signaling)
● *APC, CTNNB1* (WNT signaling)
● *BRCA1, BRCA2, PALB2* (homologous recombination deficiency)
● *MLH1, MLH3, MSH2, MSH3, MSH6, MUTYH, PMS1, PMS2, MUTYH, MBD2, MBD4, POLD1, POLE, NTHL1* (mismatch repair deficiency, see below for full characterization of MMRd)
● *AR, TP53, CHEK2, ATM, FGF4, FGFR1, HOXB13, MED12, TBL1XR1, FOXA1, CDK12, MYC, SPOP*

In general, tumor suppressor genes in the above list were required to have biallelic loss in order to be considered altered whereas known oncogenes only required a single event to be characterized as altered. This is applicable to these curated genes only. For homologous recombination deficiency genes (see above), a sample was considered to be altered if it met one of the following conditions:

● Pathogenic/likely pathogenic germline mutations (determined via Clinvar), or
● Somatic homozygous deletions, or
● Pathogenic/likely pathogenic somatic mutation (determined via Clinvar), or
● likely pathogenic somatic mutation as annotated in *Dines et 2020* ^125^or
● structural mutation, or
● gene fusion

#### Cancer Cell Fraction and mutation clonality

The cancer cell fraction (CCF) of a variant was calculated as previously described ^126^. Mutations with CCF values>0.6 were considered clonal.

#### Differential gene expression analysis in bulk RNA-sequencing

All differential expression analyses were done using the limma R package, with the default settings for the ‘‘voom’’, ‘‘lmFit,’’ ‘‘eBayes,’’ and ‘‘topTable’’ functions ^127^. Patient-level *CDKN1B* differential expression was split by disease state (localized, mCRPC) and models included coefficients for tissue site, DNA-sequencing derived sample purity, *RB1* alteration status, *TP53* alteration status, and *AR* alteration status. This allowed us to estimate log fold-changes and adjusted p-values associated while mitigating the confounding effects of tumor content, tissue site, and other prominent genomic alterations. No coefficients outside of *CDKN1B* status were used in differential expression models for LNCaP cell lines.

#### Whole genome duplication

Whole genome duplication (WGD) was calculated using ploidy estimates and the proportion of the genome with loss of heterozygosity (pLOH). Briefly, for each sample a value was calculated according to the equation: value = 3.2 - 3*(pLOH). If the sample had a ploidy both greater than this value and greater than 2.2 it was considered to be whole-genome duplicated. A similar process was used to determine samples with 2 instances of WGD. For each sample a value was calculated according to the equation: value = 5.2 - 3*(pLOH). If the sample had a ploidy both greater than this value and greater than 2.2 it was considered to have 2 instances of whole-genome duplication.

#### Chromosomal instability measures

We quantified several measures of chromosomal instability including the weighted Genome Instability Index (wGII), as well as the number of focal gains (nFG), the number of Large Scale Transitions (nLST), Telomeric Allelic Imbalances (nTAI), and Loss of Heterozygosity (nLOH) events as previously described ^13,108^. Briefly, wGII was calculated as the proportion of each chromosome that has a different copy number compared to the weighted median copy number of the sample. The average of scores for each chromosome were calculated and weighted by the length of the chromosome. For nFG, each gain had to be less than 1 megabase, have an absolute copy number value greater than 3, and be greater than the sample ploidy + 0.75. For nLST, large was defined as 10Mb and a single nLST event had a maximum distance of 3Mb between two “large segments” with allelic imbalances. For nTAI, we counted the allelic imbalances with a minimum length of 5Mb that stretched to the telomeric end of each chromosome (10Kb of chromosome start/end for GRCh38). For nLOH, we counted any segment of minimum length of 15Mb with LOH. The homologous recombination deficiency score was calculated as the sum of nTAI, nLOH, and nLST as previously described ^50,51^.

#### Recurrent copy number alterations identified using GISTIC

Recurrent copy number alterations were identified using GISTIC 2.0 ^128^. A Docker image for GISTIC 2.0.23 was obtained from https://github.com/ShixiangWang/install_GISTIC ^129^. The following parameters were used:

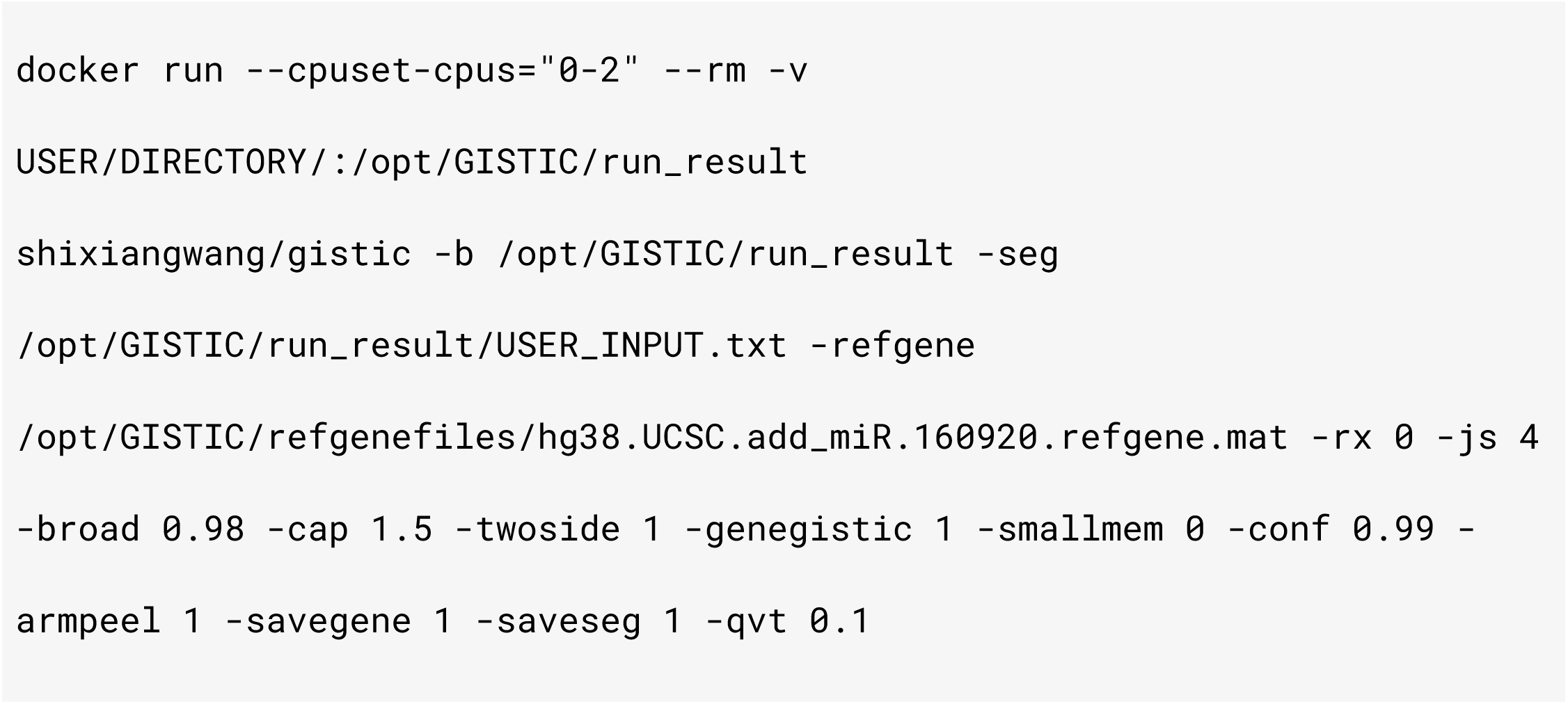

#### Copy number - gene expression correlation

To correlate the expression of genes within recurrently gained and lost regions with copy number data we utilized the following approach. First, using the all_lesions.conf_99 file provided by GISTIC, we selected all peaks with a q value < 0.01. We then utilized the amp_genes.conf_99.txt and the del_genes.conf_99.txt to select all genes within the peaks within the significant peaks. For this list of genes, we computed the ploidy-adjusted copy number (Csub-ploidy) and the reads per kilobase million (RPKM) expression for all high-quality mCRPC samples and correlated these values using a spearman correlation.

#### Survival analysis

The R packages survival and survminer were used to perform survival analyses as previously described ^130–132^. Kaplan-Meier curves and Cox Proportional Hazard models were used to compare the prognosis among variables of interest via the survfit() and coxph() functions respectively. Intervals of interest included mCRPC-free survival, overall survival from time of diagnosis, and overall survival from time of mCRPC development.

#### Pathways, gene set enrichment, and overrepresentation analysis

The Molecular Signature Database (MSigDB) was used as a source for the 1,399 gene sets comprising Hallmark, Kyoto Encyclopedia of Genes and Genomes (KEGG), C6 Oncogenic Signatures, Gene Transcription Regulation Database subset of Transcription Factor Targets (GTRD-TFT), Cancer Gene Neighborhoods (CGN), and the Human Cell Atlas immune cell signature gene sets. MSigDB gene sets were accessed using the R package Msigdbr ^133,134^. Gene Set Enrichment Analysis (GSEA) was performed in R using the R package fGSEA ^135^. We used signed -log10(p-values) from limma as input to the algorithm. Enrichment analysis for gene sets of interest was assessed using the enrichGO() function from the R package clusterProfiler ^136^. All expressed genes were used as a background list. All p-values have been adjusted for multiple-hypothesis testing using Benjamini-Hochberg correction.

#### Gene signature scoring in bulk RNA-sequencing

Scores in bulk RNA-sequencing samples were generated using the singscore R package ^137^. Briefly, Reads Per Kilobase per Million (RPKM) values are used to rank the expression of each gene within a single sample. The mean ranks are normalized to theoretical maximum values, centered on zero and summed. This allows for the scoring of a signature within a sample independently of other samples.

#### Co-enrichment analysis

Mutual exclusivity and co-enrichment analysis were performed at the sample level. The R package cooccur was used to determine significance ^138^. Briefly, the expected frequencies of co-occurrence between events based on chance are calculated and used to calculate the probability of the observed extent of cooccurrence.

#### Association of transcriptomic phenotypes with prognosis

To identify and validate cancer-relevant phenotypes linked to prognosis in lethal prostate cancer, we stratified CAPSTONE into its component Discovery and SU2C cohorts for which there were 183 and 248 available patients with a sample taken during mCRPC and high-quality sequencing. We compiled 1,399 cancer-relevant gene sets from the Molecular Signatures Database including the Hallmark, Kyoto Encyclopedia of Genes and Genomes (KEGG), C6 Oncogenic Signatures, Gene Transcription Regulation Database subset of Transcription Factor Targets (GTRD-TFT), Cancer Gene Neighborhoods (CGN), and the Human Cell Atlas immune cell signature gene sets using the R package Msigdbr ^133,134^. We filtered for gene sets with at least 4 genes profiled in our assay for a total of 1,388 gene sets.

We started by scoring each of these gene sets in the 183 patients from the Discovery cohort. We used assay-adjusted RPKM values for gene set scoring to account for variation in RNA sequencing methods (polyA vs capture) as described above. We used the singscore R package (see above) to score all gene sets ^137^. We then normalized the scores from each gene set within the Discovery cohort using the scale() function and evaluated the association with overall survival from the time of metastatic castration-resistant prostate cancer (OSmcrpc) using a Cox Proportional Hazards model. We identified 84 gene sets with a false discovery rate adjusted p-value<0.1. We ranked gene sets by significance, computed the spearman correlation among all gene set scores, and removed gene sets with R values > 0.7 and a lower rank. This reduced the 84 significant gene sets to 29 independent gene sets associated with OSmcrpc in the Discovery cohort. We then scored and scaled these 29 gene sets in the assay-adjusted RPKM values from the SU2C cohort and evaluated the association with OSmcrpc. The gene sets had to have the same direction of effects between cohorts to be considered validated. P-values in the SU2C cohort were adjusted for multiple hypothesis testing using the false discovery rate procedure.

#### Association of gene-level alterations with prognosis

To identify and validate cancer-relevant phenotypes linked to prognosis in lethal prostate cancer, we stratified CAPSTONE into its component Discovery and SU2C cohorts for which there were 183 and 248 available patients with a sample taken during mCRPC and high-quality sequencing. We integrated data from somatic mutations, gene fusions, and copy number alterations. Somatic mutations needed to pass quality control, but were allowed to be clonal or subclonal. Structural mutations were required to have an allele frequency of at least 5% in the tumor library. Gene fusions needed to meet all of the following strict quality control metrics including gmap validity, mm2 validity, HQ validity in addition to being in-frame and having more than 5 supporting reads. Copy number alterations - specifically only amplifications and homozygous deletions were considered. A select number of well-established genes including primary genetic drivers, and genes enriched in mCRPC were manually curated (see above). Calls for these manually-curated genes are available in the supplementary data. The genes were then filtered to ensure coverage of their genomic coordinates in at least 85% of the Discovery samples yielding 19,342 genes.

We then binarized each gene within each sample based on the presence of a 1 or more alteration of any type. The 19,342 genes were then filtered to have at least 2% of the Discovery cohort samples (4 patients) with an alteration yielding 3,233 genes. For each of these 3,233 genes we evaluated the association with overall survival from the time of metastatic castration-resistant prostate cancer using a Cox Proportional Hazards model. 21 genes were associated with OSmcrpc in the Discovery cohort after FDR adjustment. We then manually reduced these 21 genes down to 5 genes, as 12 genes were all adjacent on chromosome 13 near the *RB1* locus and had alterations primarily attributable to homozygous deletions consistent with known loss of function mechanisms for *RB1*. The remaining 4 genes were located on chromosome 8 in close proximity to the ADGRA2 locus but had a less significant p-value in the Discovery cohort. We then evaluated the association of alterations in these 5 genes in the SU2C cohort using cox proportional hazards model. P-values in the SU2C cohort were adjusted for multiple hypothesis testing using the false discovery rate procedure.

#### Primary genetic drivers

The following alterations were considered to be primary genetic drivers of prostate cancer - gene fusions of the ETS family of transcription factors (*ERG*, *ELK4*, *ETV1*, *ETV4*, *ETV5*, *FLI1*) ^8,139^, gene fusions of the RAF family members (*BRAF*, *RAF1*) ^9^, activating SPOP hotspot mutations ^140^, *FOXA1* class I alterations ^12^, CDK12 deficiency ^13^, germline homologous recombination deficiency (*BRCA1*, *BRCA2*, *PALB2*) ^14^, mismatch repair deficiency ^15^, and *MYC* gene fusions ^28^. Somatic HRd was not included as a primary genetic driver given the limited mutual exclusivity between somatic *BRCA2* alterations and primary genetic drivers including ETS gene fusions and *SPOP* mutations mCRPC (24,36). Mismatch repair deficiency status was computed using MSIsensor2 ^141^. A combination of copy number data, germline and somatic mutational calls, and gene expression were used to manually annotate the other primary genetic drivers. Annotation of the primary genetic driver status for each sample within CAPSTONE is present in the supplementary data.

#### Association of primary genetic drivers with likelihood of mCRPC development

To evaluate the contributions of established primary genetic drivers to mCRPC development, we compared the incidence of each driver in patients with mCRPC compared to TCGA patients with ISUP Gleason grade ≤2 localized prostate cancer. We evaluated the odds of identifying a primary genetic driver between these groups using a two-sided exact binomial test as previously described ^14^. As *CDK12*, MMRd, and *MYC* fusions were not present in any of the TCGA patients with ISUP Gleason grade ≤2 disease, inter-group odds ratios were approximated by adjusting the frequency for these alterations by 1 in each group to allow odds ratio estimation. Significance values for these groups were unchanged.

#### Primary genetic driver mutual exclusivity analysis

Primary genetic drivers including fusions (ETS, RAF, MYC), deficiencies (gHRd, CDK12, MMRd), and activating mutations (SPOP, FOXA1 class I) were evaluated across all samples. For each primary genetic driver, the samples in which the primary genetic driver were detected were evaluated for the presence of any of the other listed primary genetic drivers. The proportion of samples with only the specified primary genetic driver present were said to be mutually exclusive.

#### Identification of samples with neuroendocrine characteristics

Samples from mCRPC were clustered into 4 groups using k-means clustering, performed separately within the Discovery and SU2C cohorts. This unsupervised approach revealed a single cluster within each group that was highly enriched for neuroendocrine marker genes ^142^.

#### Generation of the CAPSTONE prognostic gene signature (CAPsig)

To nominate genes for prognostic modeling, we assembled our Discovery and SU2C cohorts consisting of 183 and 248 patients respectively. Samples exhibiting features consistent with neuroendocrine differentiation based on molecular classification signatures were excluded (see above) yielding 175 and 243 patients respectively. All expression data were batch-corrected for biopsy site effects and tumor purity. The Discovery cohort was profiled purely on the Agilent V4 transcriptomic platform. The SU2C cohort was stratified into separate cohorts by RNA profiling method - SU2C-capture included 142 Agilent V4 samples while SU2C-polyA included 101 polyA-selected RNA-seq samples. This minimized variability between cohorts attributable to known sequencing batch effects.

The nomination of prognostic genes followed a multi-stage analytical workflow conducted entirely in the Discovery cohort. Briefly, 33,063 profiled genes with low expression or low inter-sample variability were removed, leaving behind 16,531 genes. We next evaluated gene-level associations with overall survival from the time of mCRPC using univariable Cox proportional hazards regression (coxph function, R version 4.3.3), retaining 2,485 significant (p<0.05) genes. To refine the candidate gene set and define biologically meaningful patient subgroups, we performed principal component analysis on the survival-associated genes to reduce dimensionality and capture the major axes of transcriptional variation. Then k-mean clustering was used to identify two robust transcriptional clusters that reflected underlying tumor heterogeneity and distinct molecular subgroups. Limma was used to perform differential expression analysis to compare these two subgroups and identify the genes with the greatest inter-group differences. To construct a panel that was sufficiently compact for clinical implementation yet retained maximal prognostic information, we systematically evaluated nested models of increasing size (10–250 genes) and observed that performance plateaued with minimal impact beyond the top 100 genes based on fold change. On this basis, the top 100-ranked genes were selected as the CAPSTONE prognostic signature.

#### Generation of the CAPSTONE multi-omic risk classifier (CAPrisk)

Across N=431 patients (Discovery: 175 + 8 neuroendocrine, SU2C: 142 capture + 101 polyA + 5 neuroendocrine), we compiled overall survival from time of mCRPC development and mortality information along with *CDKN1B*, *TP53*, and *RB1* alteration statuses. We calculated CAPrisk scores using RPKM values adjusted for RNA profiling method (capture vs polyA) as described above. For each patient X we trained a cox proportional hazards model on N-X patients,

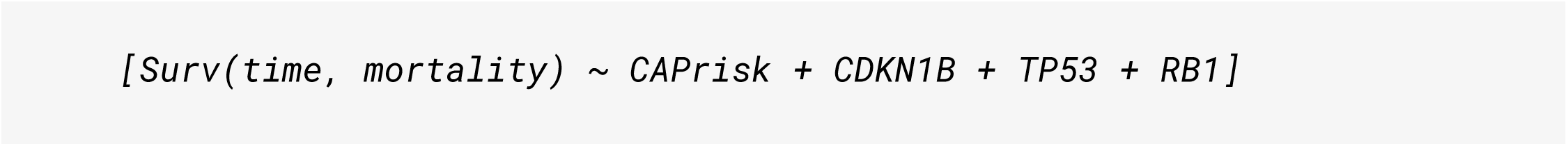

We then used the model trained on N-X patients to assign a score to the current patient X. This process was repeated for all N patients such that N total models were constructed with each patient having been assigned a score. Patient scores were then binned into 3 distinct risk groups using quartile-based stratification. Low-risk patients were in the lowest quartile of scores, intermediate-risk patients were in the middle quartiles, and high-risk patients scored in the highest quartile.

#### Evaluating outcomes in standard-of-care treatments

To identify associations between CAPrisk and standard-of-care treatments in mCRPC, we selected therapies well-represented in CAPSTONE. These included androgen receptor pathway inhibitors (ARPIs) [abiraterone, enzalutamide], taxane chemotherapy [docetaxel, cabazitaxel, paclitaxel, tesetaxel], poly (ADP-ribose) polymerase inhibitors (PARPi) [Olaparib Veliparib], and radium-223. Other therapies including Sipuleucel-T, platinum chemotherapy [carboplatin, cisplatin], and immune checkpoint blockade (ICB) [Ipilimumab, Pembrolizumab] had fewer than 30 total patients that received the therapy in an applicable context. Specifically, we only considered patients receiving each therapy that met the following criteria:

1. had a high quality sample taken during mCRPC,
2. were on the specified therapy for a minimum of 14 days,
3. this was the first time the patient had received a therapy in this therapeutic class,
4. had their mCRPC sample taken either prior to starting the therapy or within 1 year of starting the therapy, and
5. had known dates of treatment start, dates of treatment end, and dates of last follow up.

For each therapy, we focused on two intervals - Time on Therapy (ToT) and treatment-specific Overall Survival (OStxt). We evaluated the relationship between these clinical intervals and the assigned CAPrisk groups (see above) using cox proportional hazards models. Hazard ratios with 95% confidence intervals for incremental increases in risk group and unadjusted p-values were reported for ARPIs, Taxanes, and PARPi. Given the small sample size (n=35) and low frequency of use in HR patients (n=3), Radium-223, significance was assessed via log-rank test across all three risk groups.

#### CDKN1B knockdown using siRNA transfection

LNCaP (CRL-1740) were purchased from ATCC and cultured as described previously ^143^. The authenticity of the cell lines were validated with STR DNA fingerprinting by Genetica Cell Line Testing (a LabCorp brand) and regularly tested for mycoplasma contamination using the MycoAlert Mycoplasma Detection Kit (Lonza catalog LT07-318).

LNCaP cells were seeded at a density of 2.5 × 10⁵ cells per well in 6-well plates and transfected the following day using Lipofectamine RNAiMAX (Thermo Fisher Scientific Cat#13778075) according to the manufacturer’s instructions. Cells were transfected with either a non-targeting control siRNA pool (Cat#D-001810-10-50 or D-001810-10-20, Horizon Discovery) or a CDKN1B-targeting siRNA pool (Cat#L-003472-00-0005, Horizon Discovery).The siRNA oligonucleotides were transfected with Lipofectamine 3000 (ThermoFisher, cat no. L3000008) transfection reagent following manufacturer protocol for 96 h. Cell viability was measured at the endpoint using the CellTiter-Glo® 2.0 (Promega, Cat#G9243) reagent and % cell growth was normalized to siNTC. Total RNA was harvested 96 hours post-transfection and subjected to poly(A)-enriched RNA library preparation and sequencing on the Illumina NovaSeq 6000 platform as previously described ^144^.

## QUANTIFICATION AND STATISTICAL ANALYSIS

Quantification and statistical analyses were performed using R unless otherwise described. Categorical variables were compared using the chi-square test. Continuous variables were compared using Anova and Wilcoxon rank sum tests unless otherwise noted. Time to endpoint analyses were conducted using log-rank tests and cox proportional hazards models. Conventional boxplots with sample sizes are used for plotting continuous variables. Correlations between continuous variables were evaluated using the Spearman correlation coefficient. Significance thresholds were set at 0.05 unless otherwise noted. Multiple testing correction was performed using false discovery rate unless otherwise noted.

## SUPPLEMENTAL FIGURE TITLES AND LEGENDS

**Figure S1: CAPSTONE clinical endpoints and genomic characterization of disease states**

**A.** Sample treatment exposure for Discovery (DIS) and SU2C cohorts of CAPSTONE. Bars indicate the proportion of samples exposed to that therapy at time of sample collection.

**B.** Somatic homologous recombination deficiency (sHRd) pathway alteration status by disease state at time of biopsy. Chi squared test used to determine significance.

**C.** Total homologous recombination deficiency (tHRd) including somatic and germline pathway alteration status by disease state at time of biopsy. Chi squared test used to determine significance.

**D.** PI3K pathway alteration status by disease state at time of biopsy. Chi squared test used to determine significance.

**E.** WNT pathway alteration status by disease state at time of biopsy. Chi squared test used to determine significance.

**F-H.** Chromosomal instability measures by disease state at time of biopsy. Measures include **F)** number of loss of heterozygosity events, **G)** number of large-scale transitions, **H)** whole genome integrity index,

**I.** Whole genome duplication status by disease state at time of biopsy. Significance determined using Chi squared test.

**J-L.** Chromosomal instability measures by disease state at time of biopsy. Measures include **J)** number of telomeric allelic imbalances, **K)** number of focal gains, and **L)** homologous recombination deficiency score.

ANOVA and Wilcoxon rank sum test used to determine significance unless otherwise noted.

**Figure S2: Characterization of MYC gene fusions and primary genetic drivers**

**A.** MYC amplification status stratified by MYC gene fusion status and MYC amplification status.

Patients with both MYC amplification and MYC gene fusion (n=7) were omitted. Significance determined using an ANOVA and subsequent Wilcoxon rank sum tests.

**B.** MYC expression stratified by MYC gene fusion status. Patients with both MYC amplification and MYC gene fusion (n=7) were omitted. Significance determined using an ANOVA and subsequent Wilcoxon rank sum tests.

**C.** Co-occurrence or mutual exclusivity of primary genetic drivers in 1,003 prostate cancer samples.

**D.** Hazard ratio for primary genetic drivers and overall survival from time of metastatic castration-resistant prostate cancer. Point estimates represent hazard ratios, lines represent 95% confidence intervals. Arrowheads indicate the interval extends beyond range shown.

Significance: * <0.05, ** <0.01, ***<0.001.

**Figure S3: Association of pathway alterations with prognosis in lethal prostate cancer**

**A-F.** Kaplan-Meier plots for overall survival from time of metastatic castration-resistant prostate cancer (OSmcrpc) with **A)** Homologous recombination deficiency (somatic + germline), **B)** Mismatch repair deficiency (MMRd), **C)** PI3K pathway alterations, **D)** WNT pathway alterations, **E)** Cell-cycle alterations, and **F)** Cell cycle pathway alterations excluding *RB1* and *CDKN1B*. Significance determined via log-rank test.

**Figure S4: Genomic characterization of *CDKN1B* alterations in the context of neuroendocrine prostate cancer**

**A-B.** Kaplan-Meier plots for overall survival from time of metastatic castration-resistant prostate cancer (OSmcrpc) for *CDKN1B* altered patients stratified by **A)** monoallelic and biallelic loss, and **B)** homozygous deletion or mutation. Significance determined using log-rank tests.

**C.** Co-occurrence or mutual exclusivity of select genomic alterations and *CDKN1B* in 468 high-quality mCRPC samples. The significance of the relationship is represented by dot size. Abbreviations: mismatch repair deficiency (MMRd), germline homologous recombination deficiency (gHRd), FOXA1 class I mutation (FOXA1c1), FOXA1 class II mutation (FOXA1c2), FOXA1 class III mutation (FOXA1c3), somatic homologous recombination deficiency (sHRd),.

**D.** Oncoprint of 523 high-quality metastatic castration-resistant prostate cancer samples stratified by sample histology.

**E-F.** Alteration status of **E)** *RB1* and **F)** *CDKN1B* by histology.

**Figure S5: Transcriptomic characterization of samples with *CDKN1B* alterations**

**A.** Volcano plot for CDKN1B alterations in localized and metastatic castration-resistant prostate cancer samples. Differential expression models controlled for *RB1* status, *TP53* status, *AR* status, sample purity, and tissue site. Top differentially expressed genes highlighted.

**B.** Western blot for AR, CDKN1B, p53, KLK3/PSA, and histone H3 stratified by knockdown conditions.

**C.** Expression of the 10 genes represented in the NE10 gene signature in LNCaP cells with and without CDKN1B knock down.

**D.** Gene set enrichment analysis (GSEA) for the HALLMARK_ANDROGEN_RESPONSE pathway in LNCaP cells with CDKN1B knockdown relative to LNCaP cells without CDKN1B knockdown. P-value and normalized enrichment score (NES) shown.

**E.** Proliferation of LNCaP cells with and without CDKN1B knockdown. Corresponding protein levels shown.

**F.** Gene set enrichment analysis (GSEA) for the HALLMARK_EPITHELIAL_MESENCHYMAL_TRANSITION pathway in LNCaP cells with *CDKN1B* knockdown relative to LNCaP cells without *CDKN1B* knockdown.

**Figure S6: The role of *TP53* and whole-genome duplications in chromosomal instability**

**A.** Overall survival from the time of metastatic castration-resistant prostate cancer (OSmcrpc) by whole-genome duplication (WGD) status. Significance calculated using log-rank tests.

**B-E.** The B) whole-genome integrity index (wGII), C) number of large-scale transitions (nLST), D) number of telomeric allelic imbalances (nTAI), and E) number of loss of heterozygosity events (nLOH) per mCRPC sample stratified by TP53 alteration and WGD status. ANOVA and Wilcoxon tests used to determine significance.

**F.** Hazard ratio for specific recurrent gains and losses identified by GISTIC for OSmcrpc in the pooled Discovery and SU2C cohorts. Point estimates represent hazard ratios, lines represent 95% confidence intervals. Unadjusted p-values shown. Significance: * <0.05, ** <0.01, ***<0.001.

**G.** AR copy number stratified by TP53 alteration status and WGD status. AR mutation/amplification status overlaid. ANOVA and Wilcoxon tests used to determine significance.

**H-J.** CCND1, MYC, and PIK3CA alteration status by TP53 alteration status and whole-genome duplication (WGD) status in mCRPC samples. Significance determined using chi-squared test.

**K.** Androgen signaling (Hallmark Androgen Response) stratified by AR alteration status. Significance determined using ANOVA and Wilcoxon rank sum tests.

**L.** Cox proportional hazard model for OSmcrpc incorporating by TP53 alteration status, androgen signaling, and the interaction between TP53 status and androgen signaling.

**M.** Cox proportional hazard model for OSmcrpc incorporating androgen signaling. Only TP53-altered patients included.

**N.** OSmcrpc between patients stratified by *AR* alteration (mutation and amplification) and *TP53* alteration statuses. Significance determined using log-rank test. Global significance shown in plot and individual group comparisons shown adjacent to legend.

**Figure S7: Downstream characterization of the CAPsig and CAPrisk**

**A.** Gene ontology biological process enrichment for CAPsig genes. Dashed line represents adjusted p-value of 0.05.

**B.** CAPsig performance in the Discovery cohort with neuroendocrine samples included. Significance determined using log-rank test.

**C.** Significance comparison between the 20 validated transcriptomic phenotypes and CAPsig. Cox proportional hazard models with overall survival from time of metastatic castration-resistant prostate cancer development used to determine significance.

**D.** Spearman correlation between the 20 validated transcriptomic phenotypes, the prognostic gene set, and select other gene signatures.

**E.** Overlap of constituent genes between CAPsig and other gene sets including those used to create the 22-gene prognostic signature, cell cycle progression (CCP), and Basal classifiers.

**F-G.** Boxplot of Hallmark Androgen Response gene signature score for CAPrisk groups with (**F**) and without (**G**) neuroendocrine samples. Significance determined using ANOVA and Wilcoxon rank sum tests.

**H-I.** Kaplan-Meier plot for CAPrisk groups and Radium-223 outcomes for H) overall survival from treatment start, and I) time on treatment. Significance determined using log-rank tests.

## SUPPLEMENTAL TABLES

**Table S1: Sequencing quality control metrics**

**Table S2: Sample-level characteristics**

**Table S3: Clinical characteristics of 1,264 patients with prostate cancer**

**Table S4: MYC gene fusions**

**Table S5: Associations between gene-leveal alterations and overall survival from time of metastatic castration-resistant prostate cancer**

**Table S6: Assessment of redundancy for 21 gene-level associations with overall survival from time of metastatic castration-resistant prostate cancer in the Discovery cohort**

**Table S7: Associations between phenotypes and overall survival from time of metastatic castration-resistant prostate cancer**

**Table S8: *CDKN1B*-altered samples in CAPSTONE**

**Table S9: Differential expression across *CDKN1B* alterations in localized and metastatic castration-resistant prostate cancer samples**

**Table S10: Gene Set Enrichment Analysis for *CDKN1B* altered samples**

**Table S11: GISTIC recurrent gains and losses across 1,160 prostate cancer samples in CAPSTONE**

**Table S12: CAPsig constituent genes**

**Table S13: Clinical and genomic characteristics of CAPSTONE multi-omic classifier (CAPrisk) groups**

