## Supplementary figures and images for "Genomic Determinants of Lethality and Therapeutic Vulnerability in Castration-Resistant Prostate Cancer"

### Supplemental Figures S1 - S7

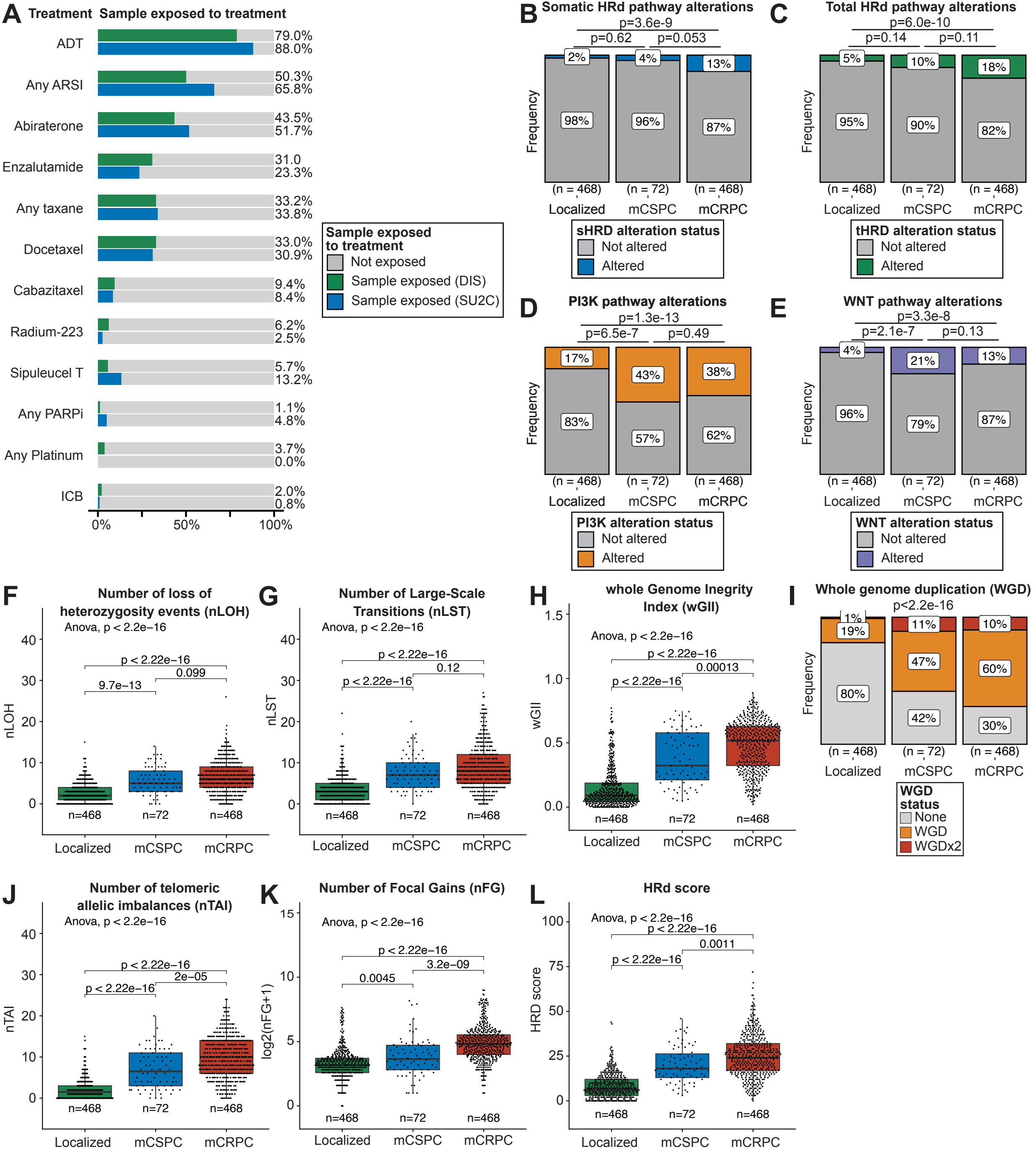

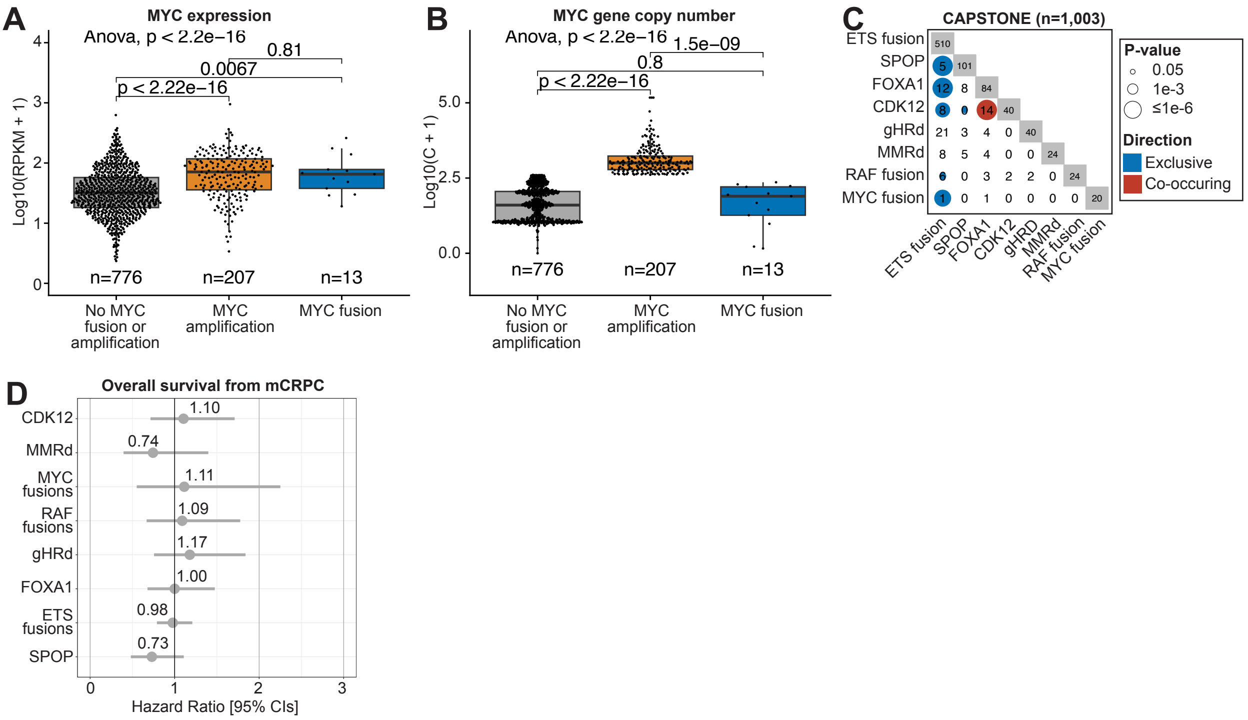

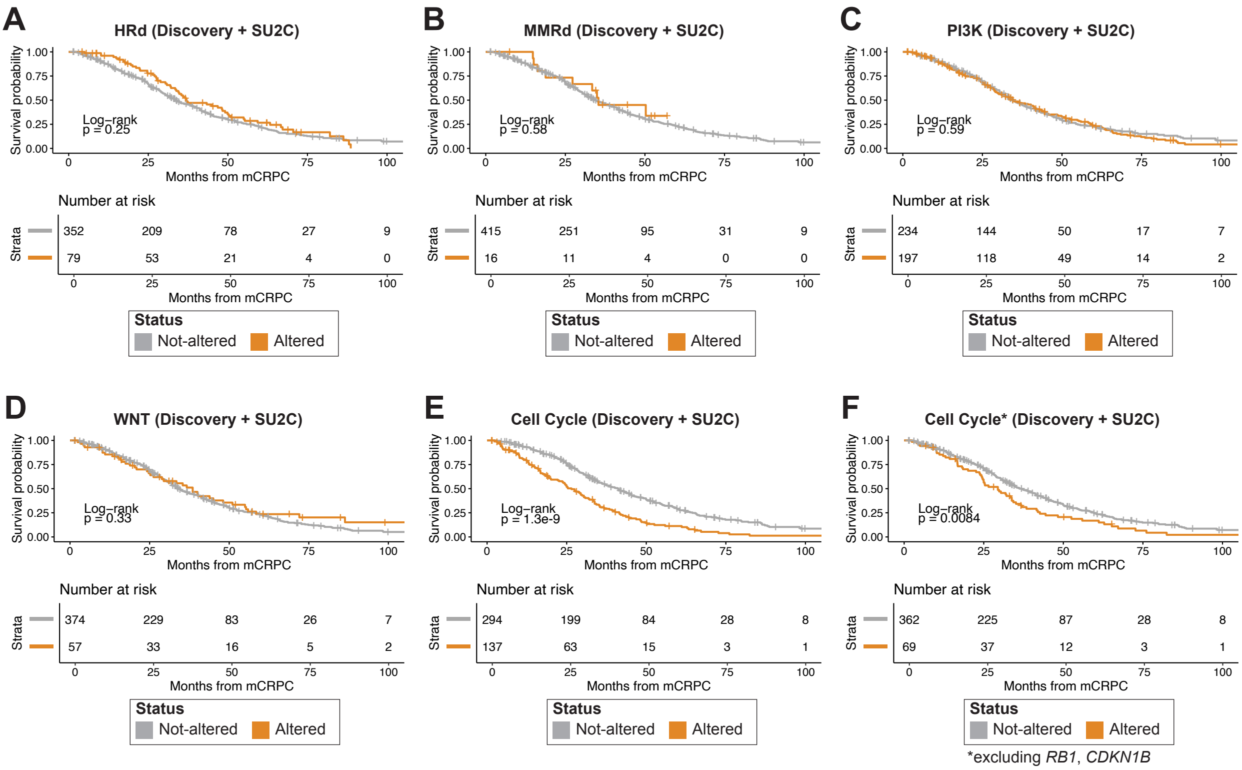

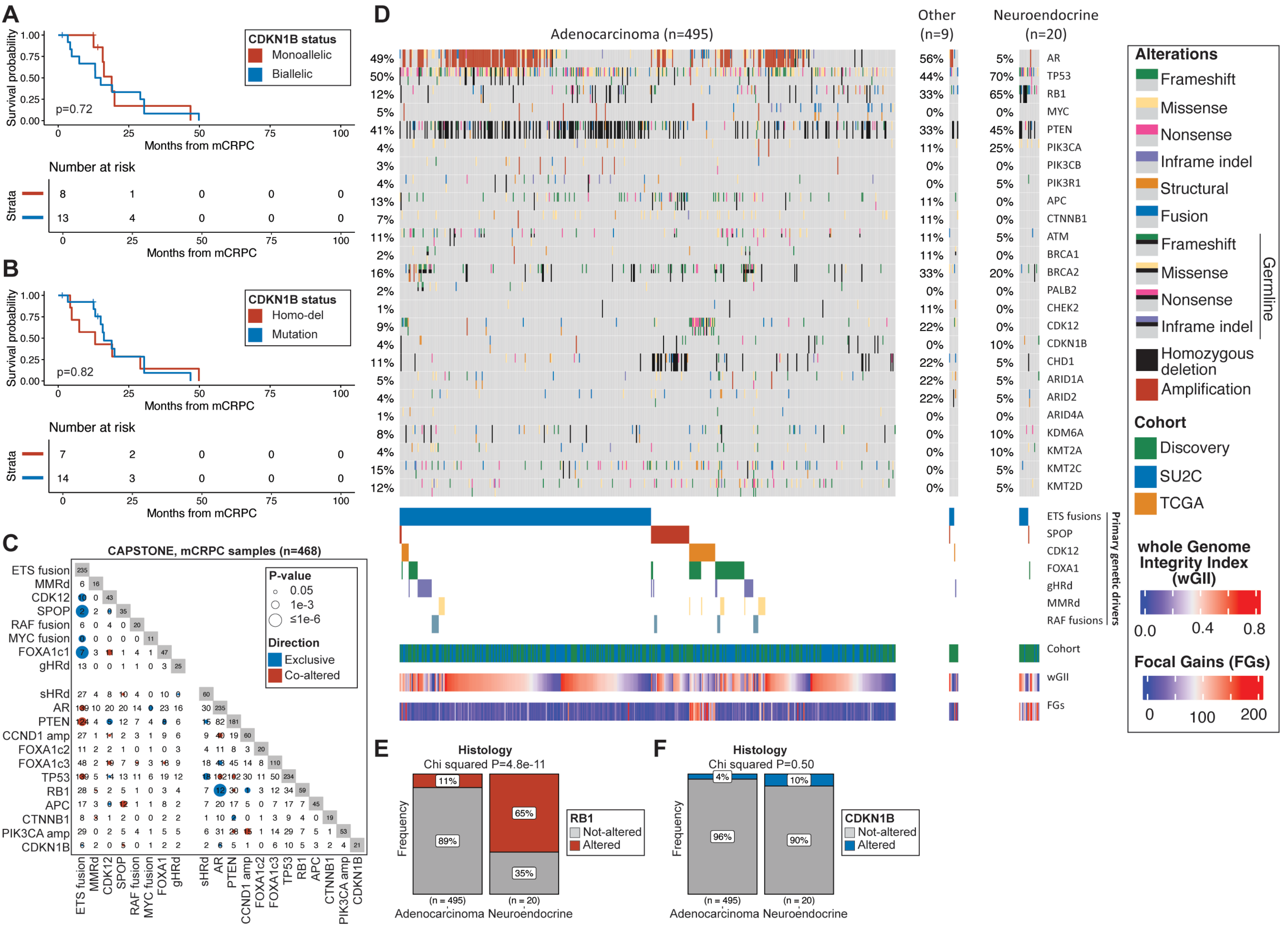

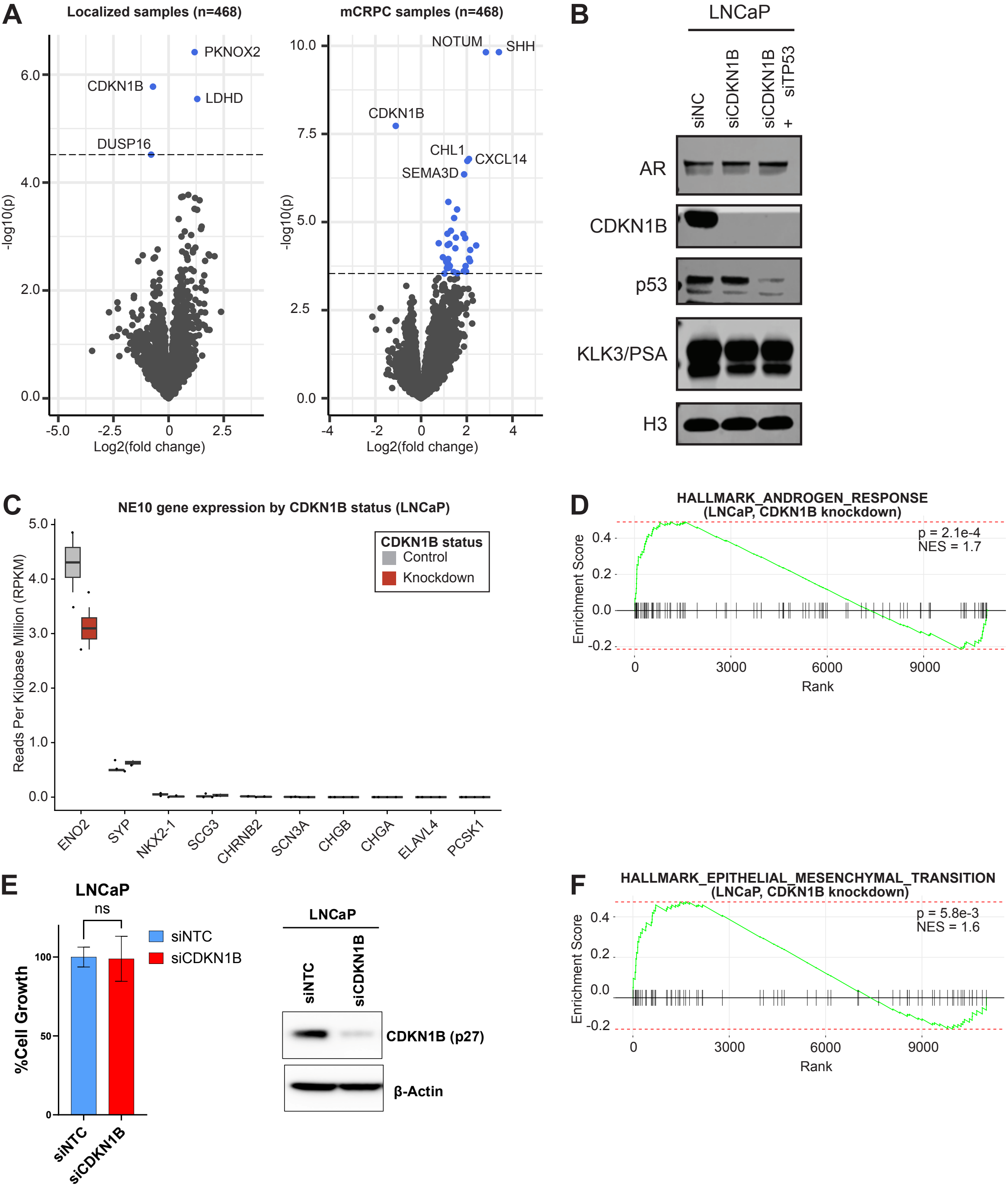

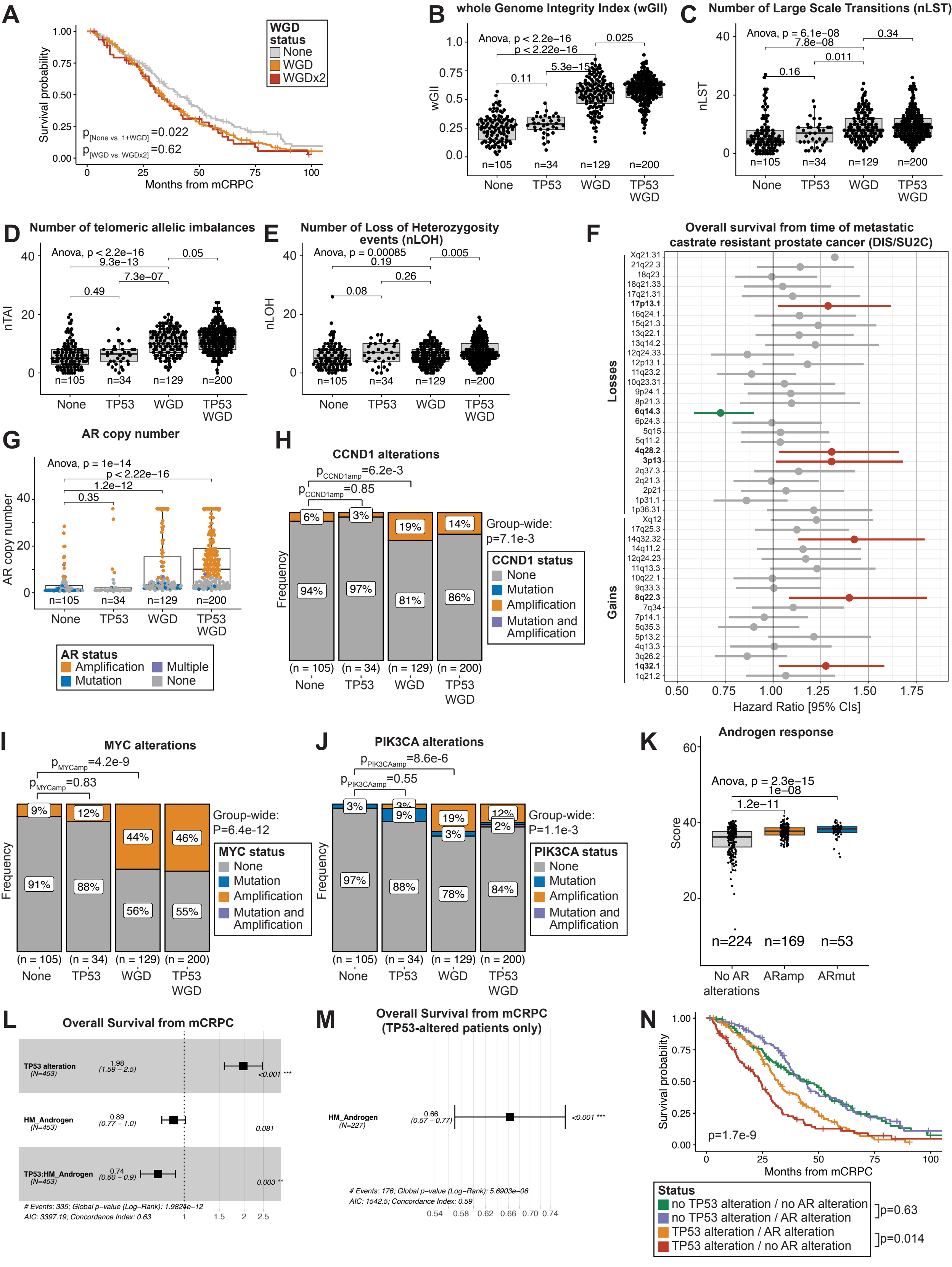

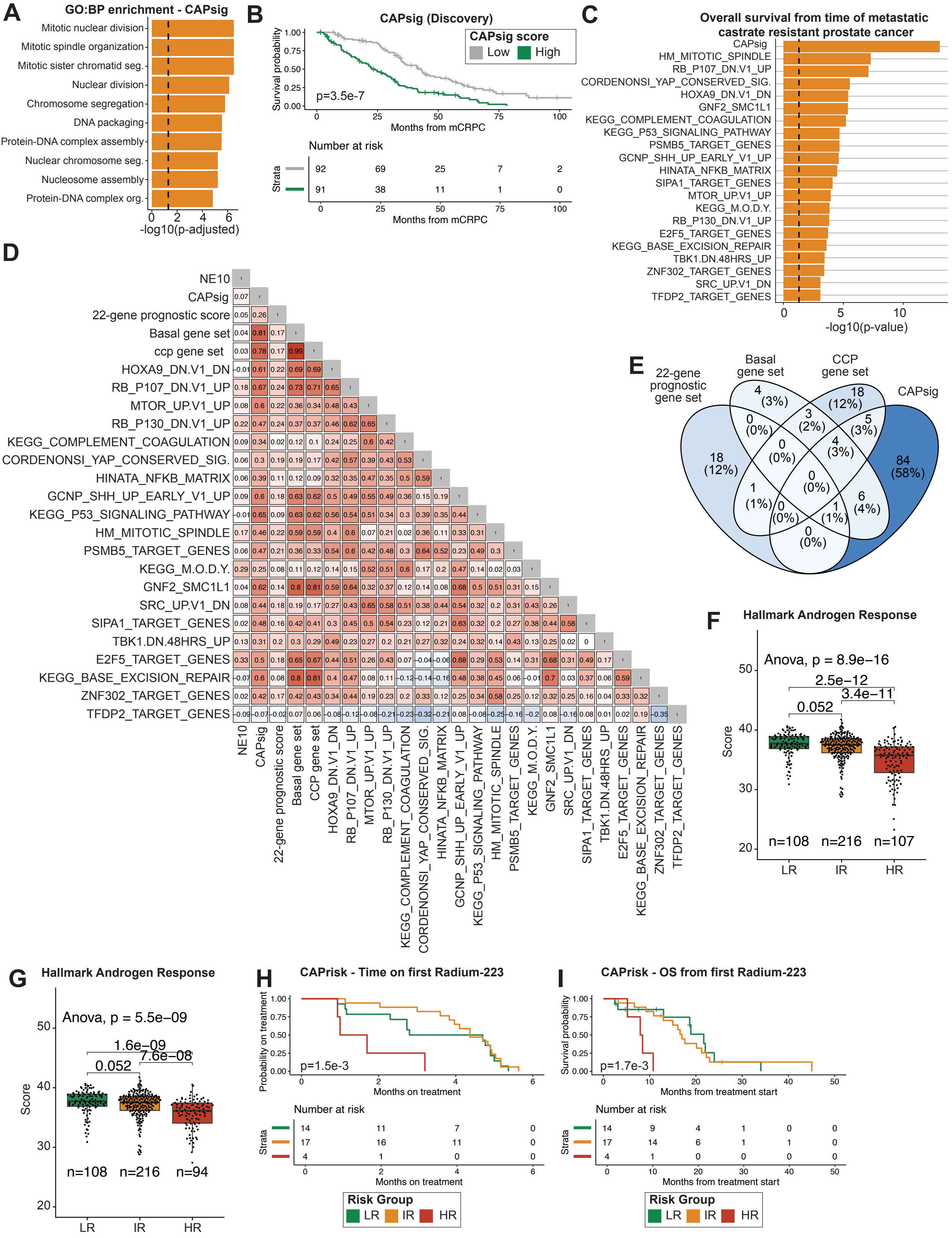
